# Development of Coupling Controlled Polymerizations by Adapter-ligation in Mate-pair Sequencing for Detection of Various Genomic Variants in One Single Assay

**DOI:** 10.1101/400689

**Authors:** Zirui Dong, Xia Zhao, Qiaoling Li, Zhenjun Yang, Yang Xi, Andrei Alexeev, Hanjie Shen, Ou Wang, Jie Ruan, Han Ren, Hanmin Wei, Xiaojuan Qi, Jiguang Li, Xiaofan Zhu, Yanyan Zhang, Peng Dai, Xiangdong Kong, Killeen Kirkconnell, Oleg Alferov, Shane Giles, Jennifer Yamtich, Bahram G. Kermani, Chao Dong, Pengjuan Liu, Zilan Mi, Wenwei Zhang, Xun Xu, Radoje Drmanac, Kwong Wai Choy, Yuan Jiang

## Abstract

The diversity of disease presentations warrants one single assay for detection and delineation of various genomic disorders. Herein, we describe a gel-free and biotin-capture-free mate-pair method through coupling Controlled Polymerizations by Adapter-Ligation (CP-AL). We first demonstrated the feasibility and ease-of-use in monitoring DNA nick-translation and primer extension by limiting the nucleotide input. By coupling these two controlled polymerizations by a reported non-conventional adapter ligation reaction 3’ branch ligation, we evidenced that CP-AL significantly increased DNA-circularization efficiency (by 4-fold) and was applicable for different sequencing methods but at a faction of current cost. Its advantages were further demonstrated by fully elimination of small-insert-contaminated (by 39.3-fold) with a ~50% increment of physical coverage, and producing uniform genome/exome coverage and the lowest chimeric rate. It achieved single-nucleotide variants detection with sensitivity and specificity up to 97.3 and 99.7%, respectively, compared with data from small-insert libraries. In addition, this method can provide a comprehensive delineation of structural rearrangements, evidenced by a potential diagnosis in a patient with oligo-atheno-terato-spermia. Moreover, it enables accurate mutation identification by integration of genomic variants from different aberration types. Overall, it provides a potential single-integrated solution for detecting various genomic variants, facilitating a genetic diagnosis in human diseases.

## Background

Human genomic variants account for ~300 Mb of sequence variation among individual human genomes and affect a variety of lengths ranging from single nucleotide to millions of base pairs^1,2^, including single nucleotide variants (SNVs), copy-number variants (CNVs) and structural variants (SVs). SNVs and CNVs are well known to be the causative factors in human diseases^3–5^, while increasing studies demonstrates the importance of detecting SVs attributed to its pathogenicity by gene-dosage change, gene(s) disruption, gene fusion or dysregulation of disease-causing gene(s)^2,6,7^. In addition, recent studies show that combination of different kinds of genomic variants in each single individual results in the variability and diversity of disease phenotypes^3,8,9^. Thus, it warrants an integrated platform for comprehensive detection of various genomic variants for each single case.

Recently, 30-fold whole-genome sequencing (WGS) with small DNA fragments (300~500 bp) has been suggested to be the first-tier testing assay for the genetic aetiological diagnosis in pediatric cases^10^. However, emerging studies have shown that at least 8.1% of those abnormalities routinely picked-up by karyotyping and chromosomal microarray analysis (CMA) would escape detection by this standard WGS^10^, letting alone those events beyond the resolutions. One of the reasons is that CNVs and SVs are predominantly mediated by repetitive elements (commonly >1 kb)^11^, causing the difficulty in identification of the flanking unique sequences by standard WGS^2,12,13^. To overcome this challenge^14^, sequencing of the junction sequences from large DNA fragments (i.e., 3~8kb) after circularization was introduced and named mate-pair sequencing^15,16^. It indicates that mate-pair sequencing with 30-fold read-depth would be an alternative strategy providing a solution to get most of the genomic variants detectable. Although there are several mate-pair methods available, limitations exist in each of them (**Table 1**), making clinical implementation impractical. For instance, the requirement of biotin labeling and enrichment results in high percentage of “inward” sequences (read pairs that do not cross the circularization junction)^15,17,18^ while random fragmentation after DNA circularization leads to a high percentage of adapter-contaminated reads (harboring adapter sequence^17^). Both of them would significantly reduce the read utility. Furthermore, a high chimeric rate exists leading to a high probability of false positives, while short-read methods^19^ reports to have a failure of detection in ~9% of cases^6^. In addition, gel-purification as a laborious procedure is mandatory in majority of methods^11,20^ (**Table 1**). Attributed to the ability of generating long reads, sequencing on third-generation sequencing platforms [i.e., PromethION (Oxford Nanopore Tech) and Sequel (PacBio, Illumina)] seems to be an alternative strategy^21,22^. However, high cost and high sequencing error rate presented prevent its application.

**Table 1.**
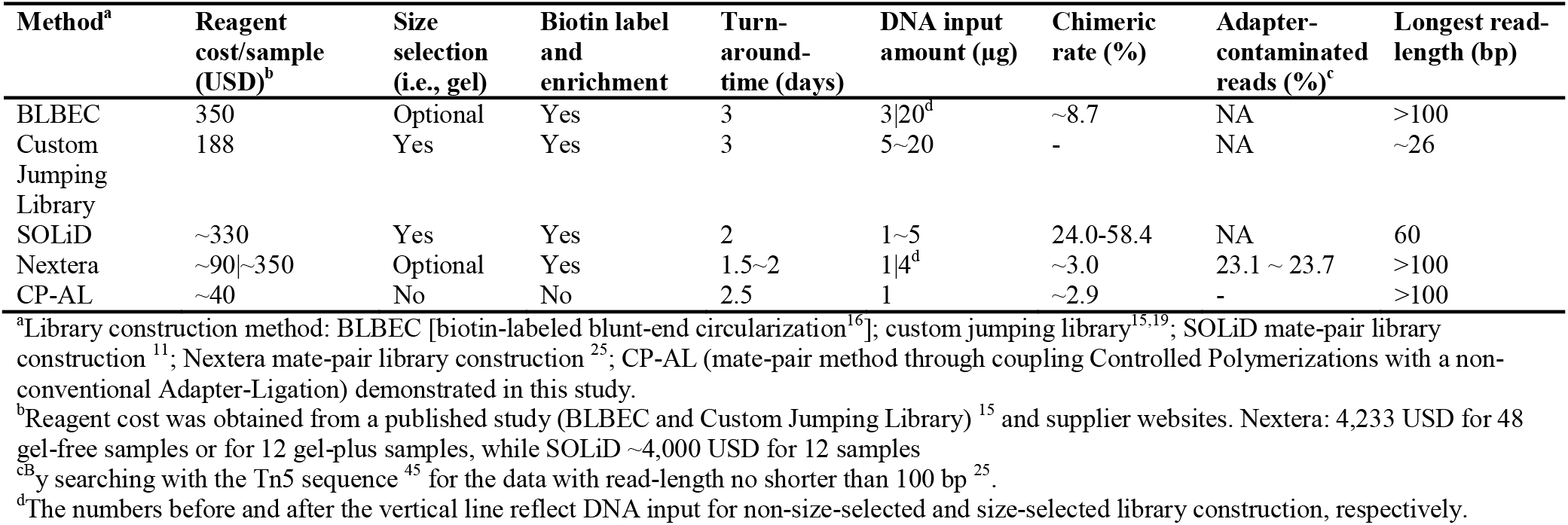
Method comparison

Recently, we discovered a non-conventional ability of T4 DNA ligase to insert 5’ phosphorylated blunt-end double-stranded DNA (dsDNA) to DNA breaks at 3’-recessive ends, gaps, or nicks to form a Y-shaped 3’-branch structure (named 3’ branch ligation or 3’BL)^23^. Given the high specificity of ligation and the achievement of 100% template usage theoretically, it provides a foundation for further development of mate-pair library construction. Herein, we describe a gel-free and biotin-capture-free mate-pair method through coupling two Controlled Polymerization reactions by this Adapter-Ligation reaction (CP-AL). Our study further demonstrates that CP-AL significantly increases circularization efficiency, fully eliminates small-inserts, and produces a narrow range of insert-size with uniform genome/exome coverage, the lowest chimeric rate and a ~50% increase of physical coverage. CP-AL were demonstrated the feasibility of being an integrated approach for detecting various genomic variants and at a fraction of current cost.

## Materials and Methods

### Ethics, consent and permissions

The study protocol was approved by the Ethics Committee of BGI-Shenzhen and The Joint Chinese University of Hong Kong-New Territories East Cluster Clinical Research Ethics Committee (CREC Ref. Nos. 2016.713 and 2017.108). Six patients with infertility or recurrent miscarriages/foetal-anomalies (detailed clinical indications were provided in **Supplementary Materials**) and with karyotyping and CMA results were enrolled, while two products of conceptions (POCs) with known trisomy were also recruited. Informed consent was obtained from each participant, ~400 μl of peripheral blood was collected in an ethylenediaminetetraacetic acid (EDTA) anticoagulant tube, while villus samples were collected from POCs. Genomic DNAs of NA19240 obtained from the Coriell Institute (Camden, NJ)^24^ was used for method evaluation, optimization and validation. In addition, published data from (1) four balanced translocations sequenced with non-size-selected biotin-labeled blunt-end circularization (BLBEC, 3~8 kb); (2) six cases prepared with the Nextera Mate-pair Library Construction Kit (hereafter called Nextera, Illumina, San Diego, CA, USA) released by the 1000 Genomes Project ^25^ were used for comparison.

### DNA Preparation and Qualification

Genomic DNA was extracted with DNeasy Blood & Tissue Kit (Cat No./ID: 69506, Qiagen, Hilden, Germany), subsequently quantified with the Quant-iT dsDNA HS Assay kit (Invitrogen, Carlsbad, CA) and QC checked. An aliquot of 1 μg DNA (OD260/OD280 > 1.8; OD260/OD230 > 2) from each sample was further sheared to a fragment size ranging from 3 to 8 kb (**Supplementary Figure S1A**) by HydroShear device (Digilab, Inc., Hopkinton, MA, USA) using the reported parameters^16^.

### Evaluation of DNA Circularization Efficiency

DNA fragments (3~8 kb) were purified with Agencourt AmpureXP beads (Beckman Coulter, Life Sciences, Indianapolis, IN, USA). After end-repair and the creation of an A-overhang, DNAs were ligated with adapter, adapter-1 (Ad1, **Figure 1** and **Supplementary Figure S2**, hereafter the adapter and primer sequences were provided in our previous study^23^). Subsequently, after annealing, to generate dsDNA with adapters incorporating dUTPs in both strands (**Supplementary Figure S3**), 320 ng of ligated DNA was amplified with a pair of primers, *Pfu* Turbo Cx (Agilent Technologies, Inc., Santa Clara, CA, USA) and extra dNTPs in a 200 μl of reaction volume according to the manufacturer’s protocol. Four micrograms of PCR products were treated at 37°C for 60 min with 10 units of UDG/EndoVIII cocktail [USER; New England Biolabs (NEB), Ipswich, MA, USA] to create a 14 bp complementary overhang. Double-stranded circles (dsCirs) were formed with one extendable gap (**Fig. 1**), and linear DNA was digested by plasmid safe (Epicentre, Illumina) at 37°C for 1.5 h. For comparison, 12 replicates were circularized by our reported BLBEC^16^ (**Supplementary Fig. S4**). Circularization efficiency was calculated as the ratio of remained DNA to the original input.

**Figure 1.**
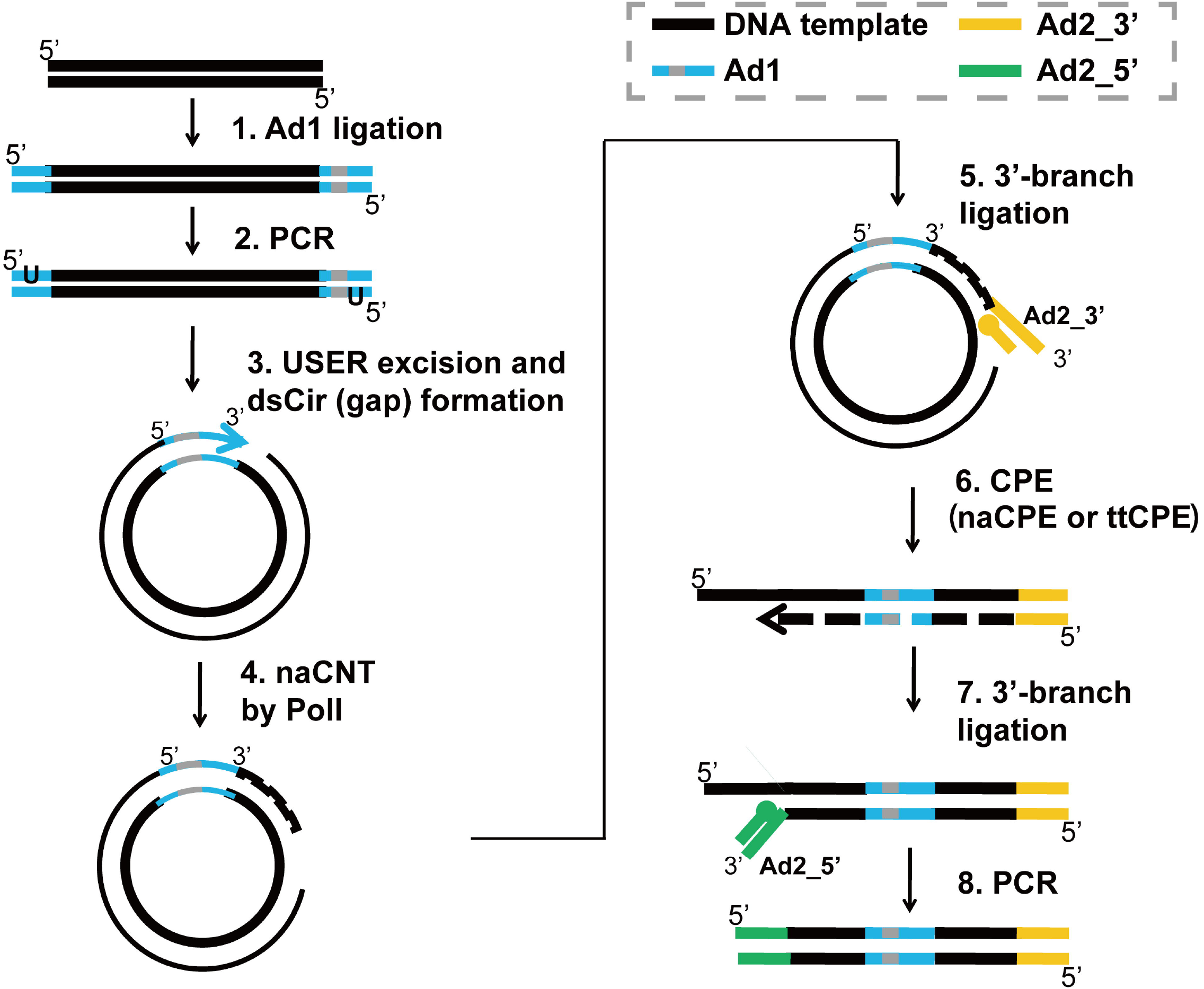
Schematic Representation of Mate-pair Library Construction by Coupling Controlled Polymerizations with a Non-Conventional Adapter-Ligation (CP-AL). The first adapter (Ad1, blue) with/without a barcode sequence (indicated by grey bar) is ligated to genomic DNA fragments (black lines). Various adapter designs may be used as the first adapter [e.g., blunt-end adapters, Y-shaped Illumina adapters, bubble adapters (**Supplementary Fig. S2**), etc.]. After Ad1 ligation and PCR, DNA ends are ligated together to form dsCirs containing a gap. naCNT is performed and results in DNA polymerization (dashed lines) and the movement of the gap into a selected length of the genomic DNA. 3’-branch ligation is used to ligate a 3’-end of the second adapter (Ad2_3’, yellow). The two strands of the dsCir are separated, and the single-stranded DNA (ssDNA) with Ad2_3’ at the 3’-end is used as a template for CPE (ntCPE or ttCPE). The Ad2_5’ sequence (green) is added to the 3’-end of the CPE product through 3’-branch ligation. After PCR, this results in genomic DNA with half of Ad2 at each end separated by Ad1.

### Evaluation of nick translation by limiting nucleotide quantities

Nick translation by PolI (DNA polymerase I) was performed in a 20 μl reaction containing 0.23 pmol dsCirs, NEBuffer 2, 10 U DNA Polymerase I (NEB), and different concentrations (10, 5, or 3.33 μM) of dNTPs (Affymetrix, Inc, Santa Clara, CA, USA) (**Supplementary Fig. S3**). The dNTP mix contained equimolar quantities of each dNTP (dATP, dTTP, dCTP and dGTP). The reaction mixture was incubated at 10°C for 20 min. The products were purified with the MinElute PCR Purification Kit (Qiagen, Hilden, Germany). Exonucleases were used for digestion and to form linear DNA^11^. Size analysis was performed for the MinElute purified Exo-treated products on 6% polyacrylamide TBE gels (Invitrogen) (**Supplementary Fig. S3**).

### Evaluation of primer extension by limiting nucleotide quantities

In brief, the 3’-ends of gDNA was ligated to the 3’-end of adapter, adapter-2 (Ad2, ON5 and ON6) and subsequently denatured at 96°C, annealed with a primer (ON10) at 56°C, and extended with *Taq* polymerase and serial dilutions of dNTPs (33.3, 25, 20, 16.7 and 14.3 pmol) at 72°C for 10 min. Products were purified with Agencourt Axygen beads and treated with Exonuclease VII, and bead purification. Size analysis was performed on a 6% TBE gel (**Supplementary Fig. S5**).

### Comparison of different parameters in CP-AL and validation

CP-AL was designed to couple nick translation and primer extension by 3’BL. For comparison, CP-ALs with different parameters (after formation of dsCir) were carried out with four DNA aliquots of NA19240^24,26^ from a single source for comparison. dsCir DNA prepared as described above was subjected to Controlled Nick Translation (CNT) and Controlled Primer Extension (CPE) in different conditions. (i) Polymerase and additional dATP and dTTP for nick translation: for Library #1, #2 and #3, 1.5 pmol of dsCirs was incubated with Polymerase I and different amounts of dNTP mix; for library #4, 0.45 U of Pol I and 0.45 U of *Taq* Polymerase were used instead; an equal quantity of each dNTP was added (33 pmol of each) for Library #1, whereas 33 pmol of dCTP and dGTP and 100 pmol of dATP and dTTP (3-fold of dCTP/dGTP in total) were added for each of the other three libraries. After nick translation, 3’BL was performed by mixing the products with the 3’-end of Ad2 (ON5 and ON6) in a 120 μl of reaction followed by bead purification; (ii) Primer extension: CPE by adjusting reaction temperature and time (ttCPE) was used for Libraries #1 and #2; the reaction contained the 3’BL product, *Pfu* Turbo Cx (Agilent Technologies), and primer ON10. The mixture was incubated at 92°C for 5 min, 56°C for 30 s and 60°C for 30 s. CPE by adjustment of nucleotide amount (naCPE) was used in Libraries #3 and #4. The reaction consisted of 3’BL products, *Taq* Polymerase, primer ON10, and different amounts of dNTPs. The naCPE program was 96°C for 5 min, 56°C for 1 min, and 72°C for 5 min. A sample of 40 μl purified product from either ttCPE or naCPE was ligated to the 5’-end of Ad2 (ON7 and ON8) and purified with Agencourt Axygen beads. (iii) Polymerase for Ad2 amplification: Libraries #1 and #4 were amplified using Q5 high-fidelity DNA polymerase (NEB). Library #2 was amplified using *Pfu* Turbo Cx. Library #3 was amplified using pyrophage polymerase (Lucigen, Middleton, WI).

For all four libraries, single-stranded circles (ssCirs) and DNA nanoballs (DNBs) were prepared and subjected to improved cPAL (Combinatorial Probe-Anchor Ligation) sequencing on a DNA nanoball (DNB)-based sequencing platform [Complete Genomics, Inc., (CGI), Mountain View, CA, USA]. We thus obtained 28 bases (2xSE28) from the 5’ ends of each adaptor, Ad1 and Ad2.^26^

### Optimized CP-AL

Purification of DNAs, end-repair, A-tailing, adapter ligation, PCR amplification (**Supplementary Fig. S1B**), purification and dsCir DNA formation were performed for each DNA, according to the procedure described above. Of note, after purification of Ad1 PCR products, a total of 20 μg amplified DNA, which could be a mixture of each amplified DNA with equal quantity (from the six patients) or from one single sample (Trisomy 2 or 8), was used for dsCir DNA formation. naCNT was performed with 1 pmol dsCir DNA; Bst DNA Polymerase, Full Length (NEB); Klenow fragment (Enzymatics, Inc.); and limited dNTPs. 3’BL was performed to ligate the 3’-end of Ad2 to the naCNT products. Primer extension was accomplished using ttCPE as described above, and the reaction mixture was incubated at 92°C for 5 min, 56°C for 60 s and 60°C for 40 s The sample was then purified with Agencourt AmpureXP beads. ttCPE products were ligated to the 5’-end of Ad2 and amplified with *Pfu* Turbo Cx (**Supplementary Fig. S1C**). ssCirs and DNBs were prepared for sequencing on the BGISEQ-500 platform (BGI-Wuhan, Wuhan, China).

For evaluation of repeatability, CP-AL was repeated for the six subjects after Ad1 PCR experiment. For these two libraries, one was subjected for sequencing with reads of 2×50 bp (equivalent to paired-end 50 bp, **Table 3, Supplementary Fig. 5B**), while the other was for 2×100 bp (equivalent to paired-end 100 bp). A minimum of 70 million read-pairs was obtained for each sample for 50 and 100 bp, respectively. For the POCs, a minimum of 30-fold read-depth was generated (2×100 bp, **Supplementary Table S3**) for each case. The robustness of CP-AL was evaluated by measurement of the DNA size for (i) DNA fragments after fragmentation with HydroShear; (ii) PCR products after Ad1 PCR and (iii) after Ad2 PCR (**Supplementary Fig. S1**).

### Data analysis, detection of genomics variants and validation

Data from CGI^26^ were aligned to the human reference genome (hg19) with a reported method^27^ and detection of SNVs and InDels were carried out by Trait-o-Matic^24,26^. For evaluation of coverage uniformity, fraction of genome/exome/coding-region was calculated as the size of region with both alleles confidently/fully called dividing by the full length (**Table 2**). In addition, sensitivity and specificity of SNV and InDel detection were determined by comparing the results from small-inserted libraries sequenced in the same platform^24^. For the data generated from HiSeq and BGISEQ (**Table 3**), alignment was performed with Burrows-Wheeler Aligner (BWA)^28^. Physical coverage was calculated as the sum of aligned distance between each uniquely aligned read-pair not resulted from PCR duplication.

**Table 2.** Evaluation of different parameters/conditions for mate-pair library construction

| Library | Nick translation | Primer extension<sup>a</sup> | % autosomal exome by GC with < 60% of mean coverage<sup>b</sup> | % fraction of the genome with fully called<sup>c</sup> | % fraction of the coding region with fully called<sup>c</sup> |
| --- | --- | --- | --- | --- | --- |
| #1 | PolI | ttCPE by *Pfu*Cx | 6.6 | 95.0 | 95.9 |
| #2 | PolI+2xAT<sup>d</sup> | ttCPE by *Pfu*Cx | 1.8 | 96.2 | 97.8 |
| #3 | PolI+2xAT<sup>d</sup> | naCPE by *Taq* | 0.7 | 96.7 | 98.1 |
| #4 | PolI/*Taq*+2xAT<sup>d</sup> | naCPE by *Taq* | 1.4 | 96.1 | 97.8 |

<sup>a</sup>ttCPE and naCPE refer to controlled primer extension by adjusting of reaction temperature and duration time or limiting the nucleotide input only
<sup>b</sup> % Autosomal exome by GC with <60% of mean coverage measures the fraction of the autosomal exome regions where the normalized coverage of exome by cumulative base GC percentage is less than 60% of the mean coverage. Higher percentage indicates higher GC bias in the autosomal exome regions.
<sup>c</sup>Fully called genome and exome fractions; both alleles can be confidently called by Trait-o-Matic software<sup>24 26</sup>.
<sup>d</sup>Additional 2-fold of dATP and dTTP relative to dGTP and dCTP

**Table 3.** Performance comparison with paired-end 50 or 26 bp reads

| Sample/data | Karyotype | Results from whole-genome sequencing | Library construction method and sequencing platform<sup>a</sup> | DNA input (μg) | Chimeric rate (%)<sup>b</sup> | Duplication rate (%) | Aligned distance <1 kb (%) | Aligned distance <1 kb (% uniquely aligned and non-duplicated) | Fraction of genome physically covered (%) |
| --- | --- | --- | --- | --- | --- | --- | --- | --- | --- |
| BCA01<sup>c</sup> | 46,XX,t(2;20)(p13;p13) | 46,XX,t(2;20)(p15;p13) | BLBEC+HS | 3 | 9.04 | 15.7 | 27.6 | 5.2 | 98.6\|91.1<sup>d</sup> |
| BCA02<sup>c</sup> | 46,XX,t(2;3)(p21;p25) | 46,XX,t(2;3)(p22.1;p26.1) | BLBEC+HS | 3 | 10.92 | 15.9 | 22.3 | 3.9 | 98.5\|93.7<sup>d</sup> |
| BCA11<sup>c</sup> | 46,XX,t(4;5;14)(q31.3;q33;q13) | 46,XX,t(4;5;14)(q32.3;q34;q21.3) | BLBEC+HS | 3 | 6.70 | 12.5 | 29.7 | 6.0 | 96.7\|91.9<sup>d</sup> |
| BCA16<sup>c</sup> | 46,XX,t(14;18)(q24;q23) | 46,XX,t(14;18)(q13.2;q22.1) | BLBEC+HS | 3 | 8.04 | 12.0 | 30.5 | 8.0 | 96.7\|91.2<sup>d</sup> |
| Sample01<sup>e</sup> | 46,XX,ins(10;13)(q11.2;q31q33) | 46,XX,ins(10;13)(q21.3;q21.2q31.3) | CP-AL+BS | 1 | 2.84 | 16.8 | 0.6 | 0.1 | 97.7\|96.6<sup>d</sup> |
| Sample02<sup>e</sup> | 46,XY,ins(6;2)(q23;p13p22) | 46,XY,ins(6;2)der(6)(6pter->6q13::6q13->6q21.3::2p16.1<-2p16.1::6q21->6q22.31::2p16.1->2p11.2::2p22.2->2p16.1::6q22.31->6q26::6q13<-6q13::6q26->6qter);der(2)(2pter->2p22.2::2p11.2->2qter) | CP-AL+BS | 1 | 2.82 | 17.8 | 0.8 | 0.2 | 96.2\|94.9<sup>d</sup> |

| Sample/data | Karyotype | Results from whole-genome sequencing | Library construction method and sequencing platform<sup>3</sup> | DNA input (μg) | Chimeric rate (%)<sup>5</sup> | Duplication rate (%) | Aligned distance <1 kb (%) | Aligned distance <1 kb (% uniquely aligned and non-duplicated) | Fraction of genome physically covered (%) |
| --- | --- | --- | --- | --- | --- | --- | --- | --- | --- |
| Sample03<sup>e</sup> | 46,XX,ins(2;18)(q31;q21.1q23) | 46,XX,ins(2;18)(q32.2;q21.1q22.1); inv(18)(q21.1) | CP-AL+BS | 1 | 3.52 | 17.5 | 0.5 | 0.1 | 98.0\|97.0<sup>d</sup> |
| Sample04<sup>e</sup> | 46,XY,der(4)ins(4)(q21.1;q31.1q31.3)inv(4)(p12q13.3) | 46,XY,der(4)(pter->p12::q13.3<-p12::q31.1->q31.23::q13.3->q31.1::q31.23->qter) | CP-AL+BS | 1 | 2.74 | 19.0 | 0.9 | 0.4 | 96.3\|95.0<sup>d</sup> |
| Sample05<sup>e</sup> | 46,XY,ins(6;3)(q13;q21q24) | 46,XY,ins(6;3)der(6)(6pter->6q14.3::3q21.1<-3q21.1::3q24<-3q21.3::3q21.1->3q21.3::6q14.3->6qter);der(3)(3pter->3q21.1::3q21.1->3q21.1::3q24->3qter) seq[hg19] del(3q24) chr3:g.146055006_148300124del | CP-AL+BS | 1 | 2.87 | 18.9 | 0.7 | 0.2 | 96.1\|95.0<sup>d</sup> |
| Sample06<sup>e</sup> | 46,XX | 46,XX | CP-AL+BS | 1 | 2.81 | 18.9 | 0.7 | 0.2 | 98.1\|97.0<sup>d</sup> |

<sup>1</sup> Sequenced with non-size-selected BLBEC libraries, published in our pilot study<sup>16</sup>.
<sup>2</sup> Sequenced with CP-AL libraries and the detection results shown here is based on the data with 100-bp
<sup>3</sup> BLBEC and CP-AL indicate library construction based on biotin-labeled blunt-end circularization and through coupling Controlled
5 Polymerizations with Adapter-Ligation, respectively; HS and BS refer to HiSeq 2000 (Illumina) and BGISEQ-500 (BGI-Wuhan), respectively.
<sup>4</sup> The numbers before and after each vertical line reflect the fraction of the genome physically coverage rate by the data with paired-end 50-bp and trimmed 26-bp reads, respectively.
<sup>5</sup> Chimeric read-pairs were defined as the read-pairs aligned to different chromosomes or to the same chromosomes but with a distance > 10 kb.

The correlation of the GC (G and C nucleotides) percentage between the human reference genome and sequencing data was assessed (**Fig. 2**). In brief, equal number (N=50 millions) of uniquely aligned read-pairs was randomly selected from each sample^29^ and merged into a dataset. Adjustable sliding windows (50 kb with 5kb increments) and non-overlapping windows (5 kb) were determined by counting an equal number of read-pairs and the reference GC percentage of each window was calculated based on the reference genome. In each case, the GC percentage of each read-pair was calculated as the reference sequence between the aligned intervals, while GC percentage of each window was set as the median value of GC percentage from all read-pairs aligned to a particular window.

**Figure 2.**
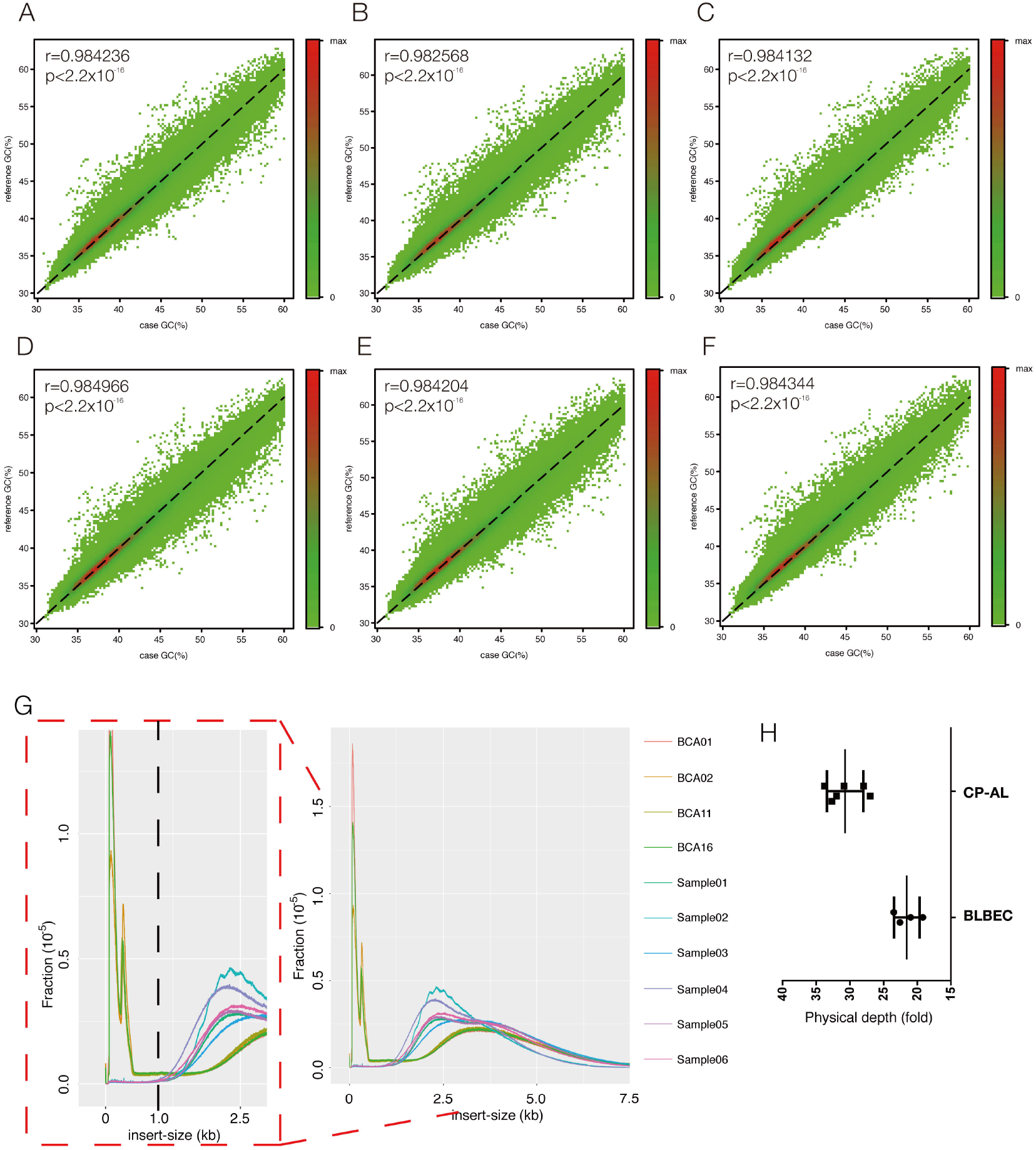
GC Concordance and Insert-size Distribution of Data from CP-AL. (A-F) The Correlation of GC Percentage between Human Reference Genome and Sequencing Reads in Sample01 to 06, respectively. The detailed method is described in Materials and methods. Each point indicates a particular window with the GC percentage (%) determined by sequencing read (X axis) and human reference genome (Y axis). Dotted line shows the ideal 100% correlation. Pearson correlation coefficient with *P*-value is shown in the left side of each figure. (G) The size distributions of four samples prepared with BLBEC and six prepared with CP-AL, respectively. Each sample was non-size-selected and sequenced with paired-end 50 bp reads. The X-axis shows the insert size, and Y-axis indicates the percentage of the paired-end reads with certain insert sizes. The figure shown in the left side with red dotted frame is the zoom in view. (H) Physical coverage distribution for the data from different library construction methods (BLBEC and CP-AL) for 50 million read-pairs per sample.

CNV analysis was performed by our published method^29^, while the SVs identification was accomplished for each sample based on a four-step procedure: event clustering, systematic error filtering, random error filtering, and aligned orientations.^13,16^ Reference sequences (1 kb) from both upstream and downstream of each breakpoint were used for primer design with Primer3 and Primer-Blast (Biotechnology Information). PCR was performed in the case and a well-established normal control (YH).^13,30^ The products were subjected to Sanger sequencing on an ABI 3730 machine (Applied Biosystems, Wilmington, DE, USA), and the results were aligned with BLAT for confirmation. Regulatory analyses such as topological associated domains and distance normalized interaction frequency were performed by the web-based 3D-genome Interaction Viewer and Database (http://www.kobic.kr/3div/).^31^ For the data from POCs, SNVs identification was carried out with Genome Analysis Toolkit HaplotypeCaller (version 3.4), and calculation of allelic ratio in chromosome-wide and genome-wide was described in the **Supplementary Methods**.

## Results

### Establishment of a new mate-pair library method: CP-AL

For clinical labs to conduct large-scale tests, an inexpensive and scalable mate-pair sequencing process is crucial. Herein, we described a new mate-pair library construction workflow through coupling two controlled polymerizations by 3’BL, which was demonstrated with higher process efficiency, lower cost, and better uniformity of genome/exome coverage in following study. In the new schema (**Fig. 1**, steps with estimated turn-around-time were described in **Supplementary Table S2**), gDNA was first fragmented into the size of interest, ligated to adapter-1 (Ad1, **Supplementary Fig. S2**), and circularized to form double-stranded circles (dsCirs). To improve the efficiency of DNA circle formation, 14 bp overhangs were used. It resulted in a median 12% circularization efficiency in 16 replicates compared with ~3% from 12 replicates from BLBEC protocol^32^ (**Supplementary Fig. S4**), a 4-fold efficiency (Wilcoxon rank-sum test: P<0.0001),

The Ad1 dsCirs contained a gap that acted as the initiation site for a controlled polymerization CNT. Subsequently, the 3’-end of adapter-2 (Ad2, Ad2_3’) was directly added to the 3’OH termini of duplex DNA containing gaps using 3’BL^23^. After ligation, another controlled polymerization CPE was performed on the single-stranded DNA (ssDNA) template. This enzymatic target enrichment strategy can efficiently eliminate the “inward” read-pairs that occur with the biotin capture method (**Table 1**). In CPE, a primer was hybridized to Ad2_3’ and extended onto the genomic template on the other side of Ad1. The 5’-end of Ad2 (Ad2_5’) was then added through another 3’BL to the newly synthesized CPE template (**Fig. 1**). Each template of the mate-pair has a selected length (an example of a minimal length of 150 bp shown in **Supplementary Fig. S6**), resulting from CNT and CPE reactions, respectively and is separated by the Ad1 sequence with Ad2_5’ and 3’ sequences at either end. The ligation products were amplified using Ad2_5’ and Ad2_3’ primers resulting in a linear dsDNA library or a ssCir DNA library that can be sequenced by various technologies such as sequencing by synthesis (SBS) on the Illumina^33^, Ion Torrent^34^ or BGI’s cPAS^35^ platforms or sequencing by ligation (SBL) using the SOLiD^36^ or first-generation CGI^26^ platforms (**Supplementary Fig. S7**).

### Feasibility testing of CP techniques for CP-AL schema

As shown previously, polymerization speed strongly depends on polymerase Km and concentration, incubation time and temperature^11^. Here we tested another hypothesis of controlling DNA extension by limiting nucleotide amount. We first used dsCirs with two gaps flanking the adapter on opposite strands as templates to assay nick translation extension (**Supplementary Fig. S3A**). Three different amounts of dNTPs (2.6, 4.0 and 8.0 pmol) were supplied in the PolI-mediated CNT reactions, which resulted in different observed extension lengths (24, 30 and 62 bp) (**Supplementary Fig. S3B**), suggesting that nick translation can be controlled by nucleotide quantity. We calculated a predicted polymerization length per arm based on the ratio of total dNTPs to the number of naCNT initiation sites (**Supplementary Fig. S3C**). For PolI-mediated reactions, observed extension lengths (24, 30 and 62 bp) were close to the predicted lengths (23, 35 and 70 bp; **Supplementary Fig. S3B and C**). It appears that PolI can utilize most, if not all, of the dNTPs in the nucleotide amount CNT (naCNT) reactions. Based on this observation, if we normalize the dsCir quantity to 1 pmol in the CP-AL schema, 100 pmol of total dNTPs will be needed in naCNT for an extension of ~100 bp. We also found that naCNT products contain gaps of a few bp, since the 3’ exonuclease activity of PolI can cleave a few nucleotides from the nicks in a 5’ to 3’ direction (data not shown).

To replace the complicated biotin capture method for targets enrichment, we tried to use CPE to recover products of interest. We hypothesize that, as for CNT reactions, there are two ways to control primer extension length—by limiting the nucleotide amount (naCPE) or by managing the reaction time and temperature (ttCPE). We first assessed whether limiting dNTPs could control DNA extension length (**Supplementary Fig. S5**). The naCPE template was 0.16 pmol of a 700~3,000 bp gDNA fraction with 3’-ends ligated to an adapter sequence. **Supplementary Figure S5B** demonstrates that different naCPE reactions generated different sizes of primer-extended products (Lane 1-5 in **Supplementary Fig. S5**), and the size correlates with the dNTP amount. Excess dNTPs (5,000 pmol, Lane 6) generated products of the original size range (700~3,000 bp). Reactions performed with the lowest quantity of dNTPs (14.3 pmol, lane 5 in **Supplementary Fig. S5**) generating the smallest products (210~300 bp). For ttCPE, the same amounts of CP-AL intermediate products after naCNT and 3’BL were extended for different times in the presence of excess dNTPs. An extension of 300~450 bp was observed after a combination of 56°C for one min and 60°C for 40 s with *Pfu* Turbo Cx. After CPE, a partially duplex DNA with a 3’ recessive end is produced. 3’BL can efficiently add adapters to gapped or 3’-recessed duplex DNA^23^. Therefore, both naCNT and CPE products are perfect templates for 3’BL.

### Comparison of different parameters in CP-AL and validation

We next compared different combinations of CP conditions and assayed the sequencing quality of the CP-AL libraries using a well-characterized sample NA19240, variants from which were reported^24,26^. To minimize the GC bias in naCNT, two approaches were used (**Table 2**). First, more than one polymerase, such as a combination of PolI with Taq Pol, was included to process through most, if not all, polymerase-specific pausing sites to minimize bias. Second, dATP and dTTP were provided in 2-fold excess relative to dCTP and dGTP (3xAT in total), given a higher percentage of A/T compared to C/G in the human reference genome.

Four libraries with different CP conditions were constructed and sequenced at a read-depth of 50~60-fold (**Table 2**). Library #1 prepared with equal amounts of four dNTPs during naCNT demonstrated the highest fraction (6.6%) of autosomal exome at <60% of the mean coverage, the lowest percentage (95.0%) of genome coverage with both alleles detected and the lowest sensitivity of SNV (95.2%) and InDel (80.7%) identification (**Supplementary Table S3**)^24^ compared to the other three libraries. It indicated that additional A/T provided during naCNT could improve the coverage uniformity. Compared with the data from small-insert libraries, with additional A/T provided, it resulted in detecting SNVs with sensitivity and specificity up to 97.3 and 99.7%, respectively, while it provided the sensitivity and specificity at 84.6 and 90.6%, respectively in InDels detection (**Supplementary Table S3**). No significant difference of genome/exome coverage was observed between Library #3 and #4, indicating no particular effect of using a combination of polymerases in naCNT. In addition, similar percentages of genome/exome coverage were observed for Library #2 (ttCPE) and Libraries #3 and #4 (naCPE), suggesting that ttCPE and naCPE were interchangeable. However, as it is difficult to precisely quantify the branched templates after naCPE and 3’BL, ttCPE was adopted. Taken together, we incorporated naCNT with 3xAT and ttCPE into CP-AL.

### Validation of CP-AL for detection of CNVs and SVs

One important application of mate-pair sequencing is CNVs and SVs detection. In this study, first, six samples (five with insertion translocations and one with normal karyotype, **Table 3**) were used for replicated CP-AL (1 μg input for fragmentation with end-products ranging from 3 to 8 kb) and sequenced for a minimum of 70 million read-pairs in 100 and 50-bp, respectively. Among these six subjects, Sample02 is a male subject diagnosed with oligo-atheno-terato-spermia, while the other five subjects have a history of recurrent miscarriages or foetal anomalies (**Supplementary Materials**). Consistent with the performance during optimization stage as described above, GC bias was minimal indicated by the significantly high concordance (r>0.98) between the GC percentage in each window calculated by human reference genome and that of the aligned reads (**Fig. 2**).

To evaluate the performance of CP-AL compared with the data generated from the most similar conditions (such as non-size-selection and 3~8 kb initial fragment size), WGS data from four samples, which were with non-size-selected BLBEC libraries and sequenced for ~50 million read-pairs (50 bp) on HiSeq 2000 platform^16^ were used for comparison. First, we compared the data utility when with the same number of read-pairs (N=50 million). Approximately 91.5% of the “inward” read-pairs in the data with BLBEC were aligned to a distance <1 kb (**Supplementary Fig. S8**), while, based on the schema design, CP-AL would not generate such read-pairs (with inconsistent aligned orientations, **Fig. 1**). Here, we compared the percentages of read-pairs with an aligned distance <1 kb. The results showed that CP-AL produced only ~0.7% of read-pairs with an aligned distance <1 kb, compared to the ~27.5% observed with BLBEC and ~24.4% with Nextera (**Table 3, Fig. 2G** and **Supplementary Fig. S9**). Approximately 33% lower physical coverage was observed using BLBEC compared to CP-AL (**Fig. 2H**), indicating a much higher utility of sequencing data using CP-AL. Of note, although DNA input was reduced from 3 to 1 μg in CP-AL, a similar duplication rate was observed in samples with CP-AL compared to those with BLBEC (**Table 3**). The reduction of PCR duplication might be owing to the increased circularization efficiency, providing more DNA templates for further experiment. In addition, to evaluate performance using shorter reads such as the data generated using EcoP15I, the read-pairs from each sample were trimmed into 26 bp in length and subjected to the analysis under the same parameters. Although no significant difference of the fraction of genome covered physically was seen between the two methods when read-depth as 50 bp, the coverage was 4% higher with CP-AL compared to BLBEC when with the 26 bp reads (**Table 3**); this increased covered fraction can be attributed primarily to the higher data utility obtained from CP-AL.

Next we evaluated the ability of CP-AL in SVs detection with different read-lengths (100, 50 and 26 bp). Compared to the results identified with 100 bp sequencing, which fine-mapped a total number of 26 rearrangements in these five insertion translocations, the diagnostic yields of data with 50 and 26 bp (trimmed from 50 bp) were only 92.3% (24/26) and 84.6% (22/26), respectively (**Fig. 3A**). It indicates that the longer read-length attained could enable a more comprehensive detection of structural rearrangements. Of note, in these five insertions the incidence of complex insertion translocations was 80.0% (4/5) (**Fig. 3B and C, Table 3**). Further, the importance of fine-mapping breakpoints can be evidenced by the potential genetic diagnosis of the patients. In Sample02, data from both 50 and 26-bp failed to identify the rearrangement in chromosome 6q13 between coordinate 74,138,669 and 74,343,934, the latter breakpoint is disrupting gene *SLC17A5* (**Figure 4A**). Although compound heterozygous mutations have been found in *SLC17A5* causing Salla disease (OMIM: #604322), no direct evidence was proved for the contribution of its heterozygous loss to spermatogenesis. However, gene *EEF1A1*, whose super enhancer was found to be highly correlated with *SLC17A5* (both genes located in the same topological domain, **Fig. 4B**)^31^, was likely haploinsufficient and might be essential for spermatogenesis^37^. It indicated that dysregulation of *EEF1A1* caused by disruption of *SLC17A5* would be the potential pathogenicity. In addition, it is also supported by the observation of disrupting 33.3% (5/15) cross-cell-type correlations between distal DNa I Hypersensitive sites and promoters of *EEF1A1* (**Supplementary Fig. S10**)^38^. However, no sample type such as RNA is available from this subject for further investigation.

**Figure 3.**
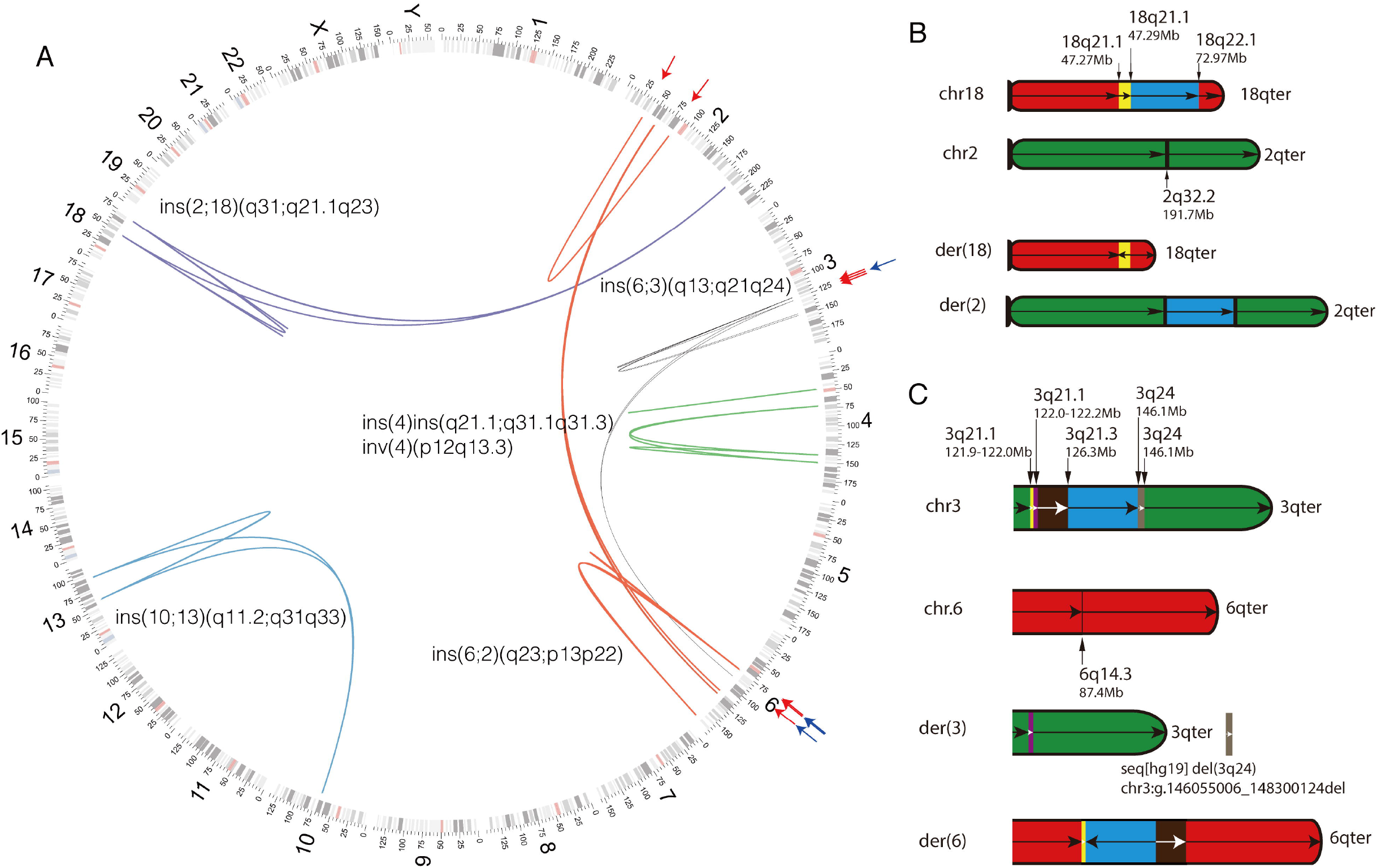
Detection of chromosomal structural rearrangements. (A) Spectrum of chromosomal structural rearrangements. Each sample is indicated with the linkages with one particular color and the event stated next to the linkages. All events are illustrated by the data from CP-AL with paired-end 100 bp sequencing and validated by Sanger sequencing. The events missed by the data from the same sample using shorter read-lengths are indicated by the red arrows (26 bp) and blue arrows (50 bp, outmost), respectively. Chromosomal nucleotide positions and bands are shown according to the University of California, Santa Cruz Genome Viewer Table Browser. The compositions of original chromosomes and the derivation chromosomes illustrated by the sequencing data from CP-AL with paired-end 100 bp reads in Sample03 and Sample05 are shown in figure (B) and (C), respectively. Each DNA/chromosome segment is shown with a different color and an arrow indicating the genomic orientation.

**Figure 4.**
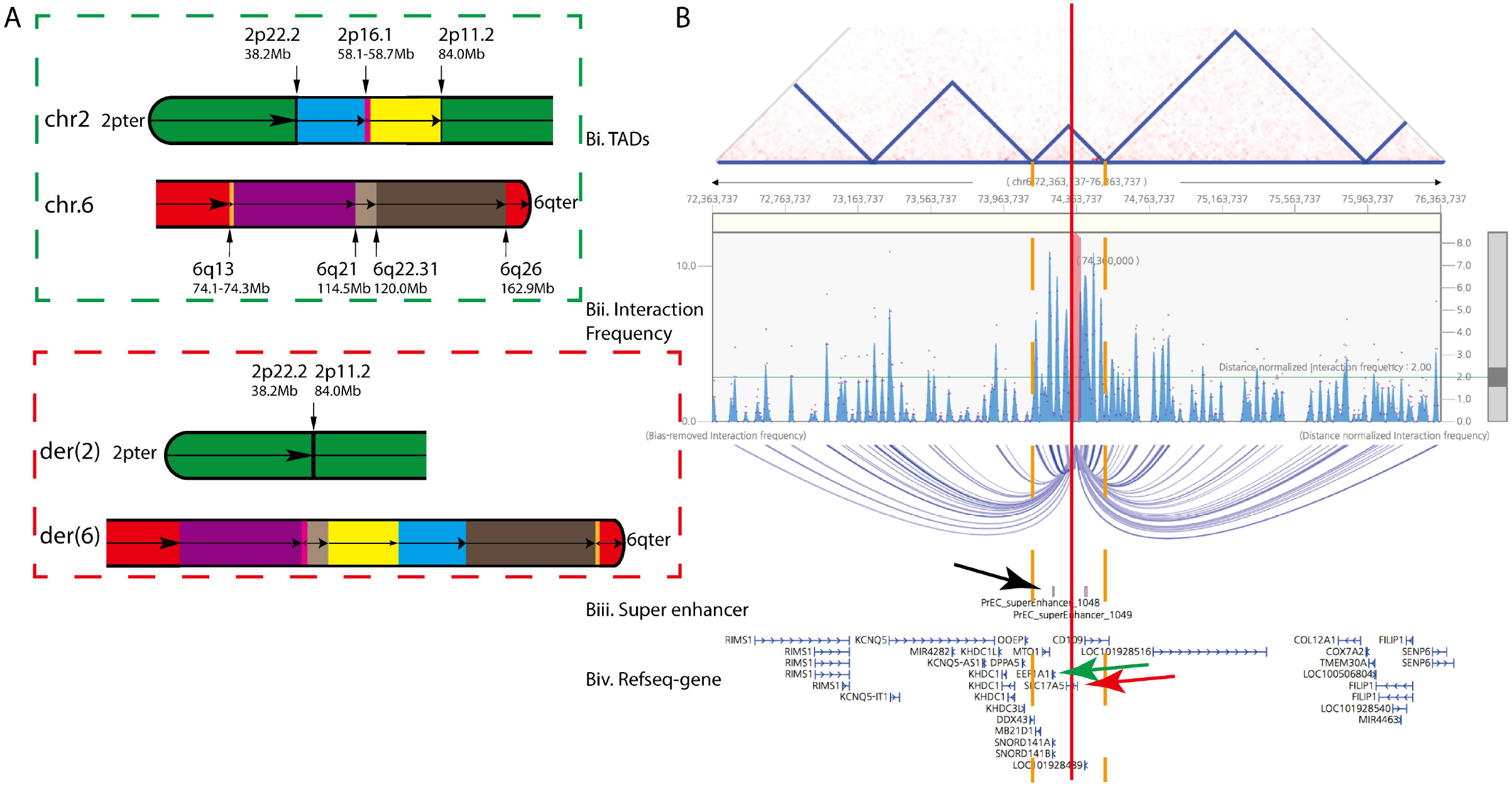
Detection of structural rearrangements in Sample02 a Male Subject with Oligo-atheno-terato-spermia. (A) The compositions of normal chromosomes (shown in green dotted frame) and the derivation chromosomes (shown in red dotted frame) illustrated by the sequencing data from CP-AL with paired-end 100 bp reads in Sample 02. Each DNA/chromosome segment is shown with a different color and an arrow indicating the genomic orientation. (B) Visualization (http://www.kobic.kr/3div/) of interaction between gene *SLC17A5* and the other locations in PrEC normal Prostate epithelial cell, BglII, the most relevant cell line in the reported database. Bi. Distribution of topological associated domains (triangles in blue); Bii. Distributions of distance normalized interaction frequency and bias-removed interaction frequency; Biii. Present of super enhancers; Biv. Distribution of genes (RefSeq). The disruption location is indicated by red line (*SLC17A5* indicated by a red arrow), while the locations of the certain topological associated domain (same as *SLC17A5*) are shown in orange dotted lines. Black arrow and green indicate the super enhancer of gene *EEF1A1* and the gene itself, respectively.

Apart from the observation of improved diagnostic yield made by the analysis with longer read-length, another advantage is to provide more uniquely aligned read-pairs in supporting rearrangement event attributed to the increased specificity of alignment with longer read-length/sequence. This was evidenced by the observation of a 33.1% reduction (from 25.1 to 16.8) of the average number of supported read-pairs for each rearrangement event in the data with 26 bp trimmed from 50-bp.

To evaluate the performance of CNV detection by CP-AL, we compared the results with CMA (**Supplementary Fig. S11**). Consistent with our previous findings, CP-AL not only reported all six rare CNVs detected by CMA, but also additionally identified 11 rare CNVs (U-test P<0.0001)^29^ including 9 deletions and 2 duplications. All of these additional CNVs were owing to the lack of probes within the targeted regions in CMA, but all of which were classified as benign or likely benign in public databases (such as ClinVar). Both assays defined a likely pathogenic deletion involving gene *ZIC1* in Sample05 (**Supplementary Fig. S12**), which was involved in the breakpoint of this complex rearrangement, indicated by the supporting read-pairs (**Fig. 3C**). Overall, our study showed a high consistency of CNV detection between CP-AL and CMA in these six cases.

### Establishment of a single assay for identification of various genomic variants

To address the possibility of identifying SNV, CNVs and SVs in one single assay, we further carried out CP-AL with >30-fold read-depth WGS for two POCs (**Fig. 5** and **Supplementary Fig. S13**). Detection of SNVs, CNVs and SVs were performed independently and integrated afterwards. In case Trisomy 2, the increased allelic ratio (**Supplementary Materials**) in chromosome 2 confirmed that there was one copy gain of whole chromosome 2 consistently with the result from CNV analysis, while SV analysis provides the information of the duplication/deletions supporting read-pairs (**Fig. 5**) for narrow-downing the critical affected regions (**Supplementary Figure S14**). Of note, the breakpoints of these deletions/duplications were predicted to locate in the regions fully with repetitive elements and segmental duplications (**Supplementary Fig. S14**), which was unlikely detectable by standard WGS. The feasibility of detecting the same genomic variants by different analyses demonstrated an accurate detection, emphasizing the advantage of integration of all detection results. A similar finding was observed in case Trisomy 8 (**Supplementary Fig. S13**), demonstrating the robustness of this application. In addition, in case Trisomy 2, a 51.4 kb inversion was detected in chromosome 6q14.1, while a 187.9 kb inversion was observed in chromosome Xq28. Consistent with the findings of deletions/duplications, the breakpoints of both inversions were also predicted to locate in the regions densely populated with repetitive elements and segmental duplications (**Supplementary Fig. S14**). Overall, the analysis from these two samples emphasizes the advantages and superiorities of applying CP-AL with 30-fold WGS as an integrated method for potentially comprehensive study of genomic variants in human diseases, compared with standard WGS.

**Figure 5.**
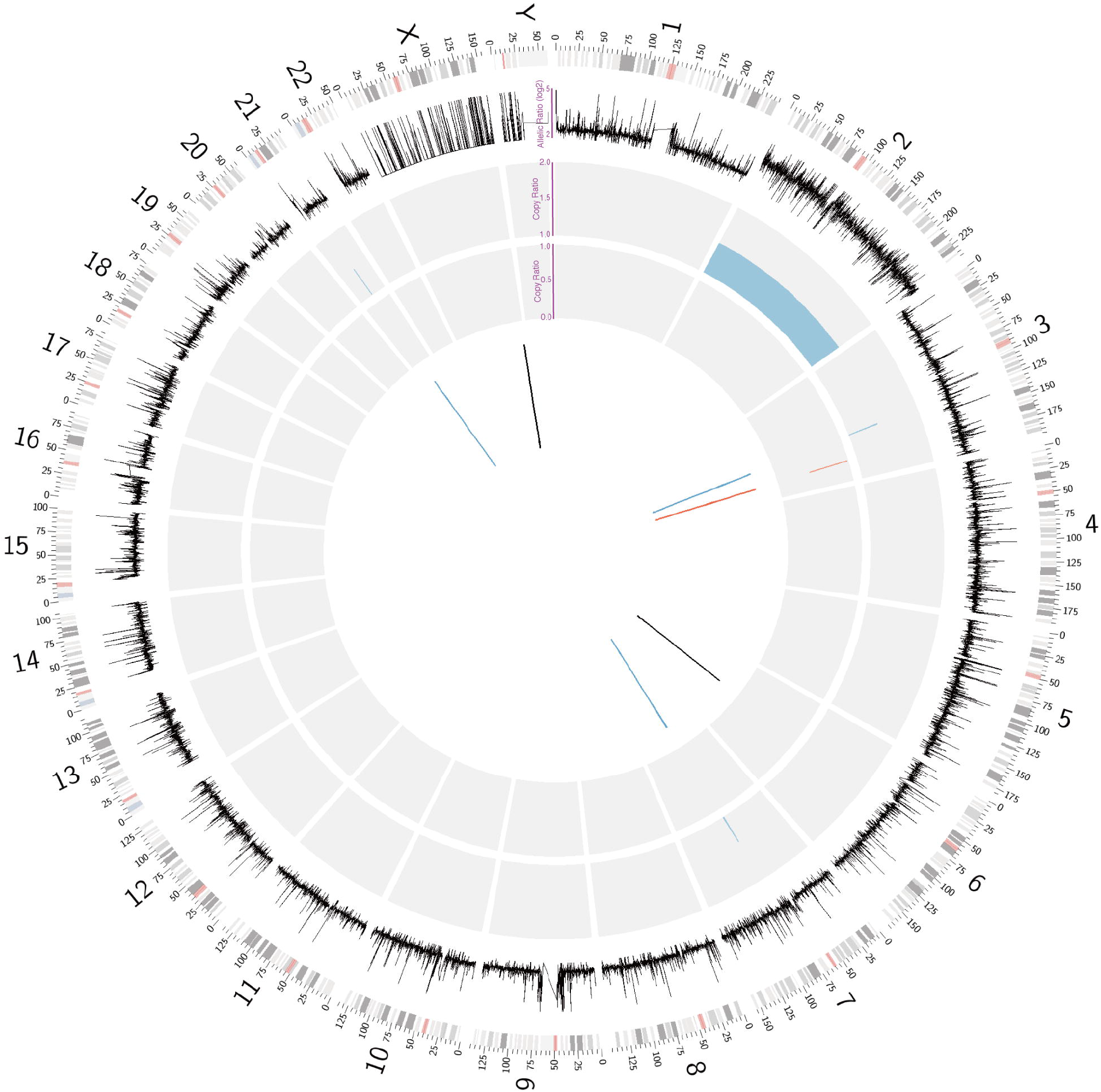
Whole-genome analysis of genomic variants in sample with Trisomy 2. The distributions of allelic ratio (window-size: 100 kb in log2 scale; **Supplementary Materials**), copy-ratio and structural variants are shown from outer to inner circles accordingly. Allelic ratio of on average 4 is shown in chromosome 2 indicating there are two copies of one base type against only one copy of another base type. Karyotypical structures and cytogenetic band colors are shown according to the University of California, Santa Cruz Genome Viewer Table Browser and chromosome color schemes (outmost circle). Rectangles in red and blue indicate copy-number losses and gains, respectively, while lines in red and blue also indicate copy-number losses and grains. Lines in black in SV analysis show two inversions detected in chromosome 6 and X, respectively.

## Discussion

Mate-pair sequencing provides a foundation for identifying various genomic variants, including SNVs, CNVs and SVs in one single assay, however, problems such as low data utility and high cost limit the clinical application of current methods (**Table 1**)^11,15^. In this study, we developed a new mate-pair method CP-AL through coupling controlled polymerizations with a non-conventional adapter-ligation reaction recently developed in our previous study^23^, and further demonstrated that its feasibility and advantages in detection and delineation of various genomic variants in human diseases.

In this study, based on the foundations provided by 3’BL reaction, we have demonstrated the effectiveness of controlling polymerizations during nick translation and primer extension and coupling these two reactions by 3’BL. Before building up the whole workflow, we demonstrated the feasibility of carrying out each key component, respectively: (i) conducting nick translation by adjusting the dNTP quantities (naCNT) instead of adjusting reaction temperature and duration time, the latter commonly results in a large size range of templates^11^; (ii) performing primer extension by either adjusting the dNTP amount (naCPE) or adjusting reaction temperature and duration time (ttCPE). Further, during optimization, a more uniform genome/exome coverage was achieved by providing a 2-fold excess of dATPs/dTTPs in nick translation: the percentage of autosomal exome with <60% of mean coverage was improved from 6.6% to 0.7%, whereas percentages of coding regions with both alleles detected in a well-characterized sample NA19240 were increased from 95.9% to 98.1% (**Table 2**).

Compared with the reported approaches^6,11,15,16,39,40^, CP-AL provides the following advantages: (i) an adapter with a 14 bp overhang is used increasing circularization efficiency (from ~3% to ~12% by 4-fold), thus decreasing the minimal DNA input amount; (ii) no gel-purification; (iii) biotin-capture-free fully eliminates the “inward-sequences” (0.7%, or a 39.3-fold reduction from 27.5% for BLBEC; 24.4% for Nextera); (iv) secondary-fragmentation-free avoids the random presence of “adapter-contaminated” (~23% from Nextera) due to random fragmentation after DNA circularization; (v) sequencing with different read-lengths and different platforms demonstrates a more uniform genome/exome coverage attributed by minimal GC biases^41^; and (vi) generation of a long read-length (100 bp or even longer) with the lowest chimeric rate provides the potential foundation for disease precision diagnosis and disease-causing gene discovery. It was evidenced by the observation of an 18.2% increased detection rate for SVs (4/22, four additional events detected from 22 detected with 26 bp), one of which potentially causes a positional effect for male infertility. In evaluation of the performances other than the consistency of variants detection^16^, it is important to have the same DNAs as input, however, no such material was available from the previous study for CP-AL experiments. Given the targeted performances were only related to the critical DNA fragment size instead of the certain mutations/variants pattern in a particular case and a consistent fragment size of the DNA input after fragmentation (**Supplementary Fig. S1A**), our study is able to support the argument of CP-AL is superior to the existing mate-pair method in these related applications.

In addition, our data further suggests that good performance of SNV and InDel detection in CP-AL (**Supplementary Table S3**). Together, CP-AL with WGS has been proved to be the most integrated assay in providing the detection results of point mutations (SNVs), deletions/duplications (CNVs), translocations, inversions and insertions (SVs). In addition, we further demonstrated its advantages by showing the higher detection accuracy attributed by detecting the same variants by different types of analyzing methods (**Fig. 5**). For example, increased allelic ratio from SNV detection confirmed the copy-number gain while SV analysis provided the read-pairs supporting the deletions/duplications (**Fig. 5** and **Supplementary Fig. S13**). Since the breakpoints of those SVs identified by CP-AL are predicted to locate in the genomic regions densely populated with repetitive elements and segmental duplications, which are unlikely detectable by standard WGS, our study shows the potential of replacing standard WGS by CP-AL.

Apart from high read-depth application, we also show the feasibility of using CP-AL with low-pass (or low-coverage) WGS for CNV and SV detection application, given that an ~50% increased physical coverage obtained from CP-AL. In this aim, only a limited number of read-pairs (50-70 million) is required in each sample; a pooling step is suitable^16^. In addition, due to the flexibility of sequencing orientation for ssDNA, pooling was performed soon after Ad1 PCR (**Fig. 1**). This change will reduce the labor cost for all subsequent procedures by treating the pooled library as a single sample. By modification of the adapter sequence, our study also shows the potential of wider applications in different sequencing platforms (**Supplementary Fig. S6**). For the cost aspect, avoiding the usage of expensive reagents such as biotin and exonuclease (exonuclease digestion and/or streptavidin beads is costly and inefficient), the reagent cost of CP-AL is only a fraction of current cost, for instance: it is only ~1/10^th^ that of BLBEC (US $40 vs $350, **Table 1**)^15^. It is important that reduction of mate-pair library cost can be achieved, given the era of 1,000 USD for standard WGS came, CP-AL would overcome the main challenges in detecting various genomic variants but with similar cost.

Moreover, the biologic interpretation of the genomic variants is important after identification. Five cases with insertion translocations known to result in abnormal embryos/pregnancies with high risk (**Supplementary Fig. S15**) were studied in this study. CP-AL with 100 bp was able to detect and delineate all the rearrangements, providing a much higher incidence of complex insertion translocations as 80% (4/5) (**Fig. 3B and C**) compared with the previous study^42^. The importance of fine-mapping the related breakpoints was demonstrated in providing the potential foundation for disease precision diagnosis and disease-causing gene discovery, such as showing a potential positional effect in a patient with oligo-atheno-terato-spermia (Sample02, **Fig. 4**). In addition, in Sample05 with cryptic deletion (not detected by karyotyping) involving OMIM disease-causing gene *ZIC1* likely haploinsufficient (HI=2) defined by ClinGen Genome Dosage Map. No clinical recognized malformation was identified in this patient. It might be due to incomplete penetrance, variable expressivity or the potential pathogenicity of this gene was contributed by gain of function instead^43^, warranting a comprehensive clinical assessment.

Although reduction of both labor work and reagent cost makes CP-AL more practical for routine practice, several limitations remain: (i) PCR amplification with primers incorporating dUTPs is used, which would be inefficient for DNA >10 kb. Foreseeing that CP-AL with limited ability in identification of SVs mediated by low-copy repeat, application of third-generation sequencing platforms^21,22^ might be an alternative strategy; (ii) Read-length: given that improvement of variants detection and delineation were observed by longer read-length attributed to the improvement of alignment specificity (**Fig. 3A**), it is valuable to provide longer read-depth (>150 bp or 300 bp, **Supplementary Fig. S6**) for CP-AL; and (iii) DNA samples were fragmented physically (Hydroshear); alternative methods such as fragmentation by Transposases (i.e., Tn5)^44^ is warranted.

In conclusion, we provide a cost-effective and high-efficiency mate-pair method CP-AL that offers higher utility of read-pairs, better uniformity of genome/exome coverage at a fraction of current cost. With 30-fold read-depth, this approach shows promise as an integrated method and eligible first-tier WGS testing assay for detection and delineation of various genomic variants in human diseases.

## Availability

Whole-genome sequencing data used in this study has been made available in the CNGB Nucleotide Sequence Archive (CNSA: https://db.cngb.org/cnsa) under the accession number CNP0000078.

## Acknowledgements

We would like to acknowledge the ongoing contributions and support of all Complete Genomics and BGI-Shenzhen employees, in particular the many highly skilled individuals that work in the libraries, reagents, and sequencing groups that make it possible to generate high-quality whole-genome sequencing data. We thank Dr. James Lupski, Dr. Sau Wai Cheung, Dr. Ya Gao and Dr. Pengfei Liu for helpful discussions surrounding manuscript preparation. Zirui Dong thanks the support of the Dr. Stanley Ho Medical Development Foundation.

