## Supplementary Materials and Methods for "Development of Coupling Controlled Polymerizations by Adapter-ligation in Mate-pair Sequencing for Detection of Various Genomic Variants in One Single Assay"

**Sample enrollment and clinical assessment**

Informed consent was obtained from each participant and detailed clinical information was recorded for further analysis. Among six subjects collected, Sample02 was a 29-year-old male patient presenting with a history of one year of primary infertility while the other five subjects were reported to have a history of recurrent miscarriage or fetal anomalies. The detailed clinical information for each sample was provided as below:

Sample01, female, G2P1, had a clinical indication of having a history of a stillbirth occurred at 37 gestational weeks. No clinical malformation was found in the patient and the husband. G-banded chromosome analysis reported an apparently inter-chromosomal insertion translocation 46,XX,ins(10;13)(q11.2;q31q33).

For Sample02, semen analysis revealed severe oligozoospermia (0.9 million/ml) with only 37.5% sperm motility (>50% in normal male) and >90% abnormally shaped sperms (<50% in normal male). Ultrasound evaluation of prostate showed enlarged size and calcification. There were no other abnormalities, medical or family history in the patient and his female partner. G-banded chromosome analysis reported an apparently insertion translocation 46,XY,ins(6;2)(q23;p13p22).

Sample03, female, G4P1, had a female child with intellectual disability in 2^nd^ pregnancy and suffered consecutive recurrent first-trimester miscarriages in her 3^rd^ and 4^th^ pregnancies. No clinical malformation was detected in the patient and G-banded chromosome analysis reported apparently insertion translocation 46,XX,ins(2;18)(q31;q21.1q23).

Sample04, male, with his wife had previous child/pregnancy reported with chromosome abnormality: arr[hg19] 4q31.1q31.23(140046328_150534134)x1 in the product of conception (POC, **Supplementary Figure S15**). G-banded chromosome analysis of the patient reported an apparently intra-chromosomal insertion translocation involving an pericentric inversion in the breakpoint regions 46,XY,der(4)ins(4)(q21.1q31.1q31.3)inv(4)(p12q13.3). No clinical malformation was found in the patient and the wife.

Sample05, male, G3P1, with his wife had a stillbirth in 1^st^ pregnancy, a male child diagnosed with chromosomal abnormality, and a fetus with ultrasound structural anomalies. G-banded chromosome analysis in this patient reported an apparently inter-chromosomal insertion translocation 46,XY,ins(6;3)(q13;q21q24). No clinical malformation was found in the patient and his wife. CMA reported a likely pathogenic deletion arr[hg19] 3q24(146172283_148219238)x1 (**Supplementary Figure S12**) in this patient.

Sample06, female, had a history of recurrent miscarriages and G-banded chromosome analysis reported a normal karyotype. No clinical malformation was found in the patient and the husband.

**Chromosomal Microarray Analysis**

CMA was performed in each of the six patients by using a well-established customized 44K Fetal DNA Chip v1.0 (Agilent Technologies, Inc., Santa Clara, CA) according to the manufacturers’ protocols and the data were analyzed via CytoGenomics^12^.

**Genome-wide allelic ratio calculation**

For whole-genome sequencing (WGS) data (>30-fold) from those two cases with Trisomy, we further evaluated the feasibility of identification of copy-number variants (CNVs) by utilizing the single-nucleotide variants (SNVs) data.

For duplication analysis, allelic ratio from each SNV was defined as ratio of the larger number of allele supporting reads dividing by the smaller. Since for the heterozygous SNVs, the average value of allelic ratio is 1.63 while the standard deviation is 1.30, we further set allelic ratio as 3.0 as the cutoff for selecting the heterozygous SNVs for further analysis. In order to reduce the deviation, 100-kb fixed window size was used and the average allelic ratio within each window was used as the allelic ratio for this particular window. For those two cases with CP-AL library construction and 30-fold WGS read-depth, we observed an increased allelic ratio in chromosome 2 (**Figure 5**) and chromosome 8 (**Supplementary Figure S13**) supporting Trisomy 2 and Trisomy 8, respectively.

For deletion analysis, 100-kb window size was used and the ratio of the number of heterozygous SNVs against the number of homozygous SNVs was calculated and used for consideration of deletion or loss of heterozygosity (data not shown).

**Supplementary Figures**

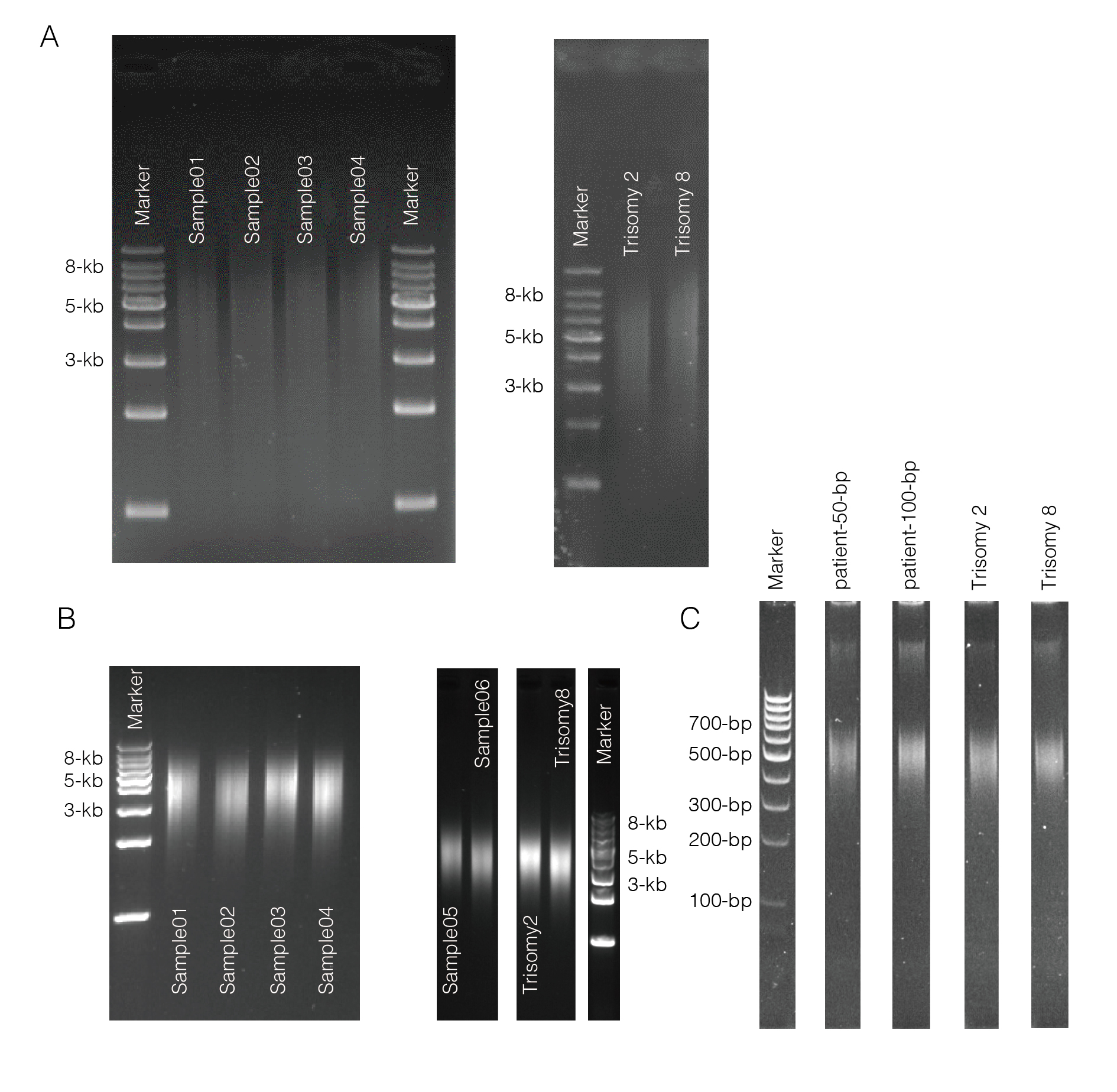

**Supplementary Figure S1.** **Size distribution of DNA fragments before CP-AL, after adapter-1 PCR and after adapter-2 PCR**. (A) Gel figures show the size distribution of DNA fragments after DNA fragmentation by the HydroShear device with the same parameters reported in our pilot study^3^. Gel figures indicate the main size of each sample is ranging from 3~8-kb with the dominant size in 5-kb. (B) Size distribution of DNA fragments after adapter-1 PCR experiments. (C) Size distribution of the final products for sequencing (after adapter-2 PCR). Gel figures show each library has a confined size 400~600-bp.

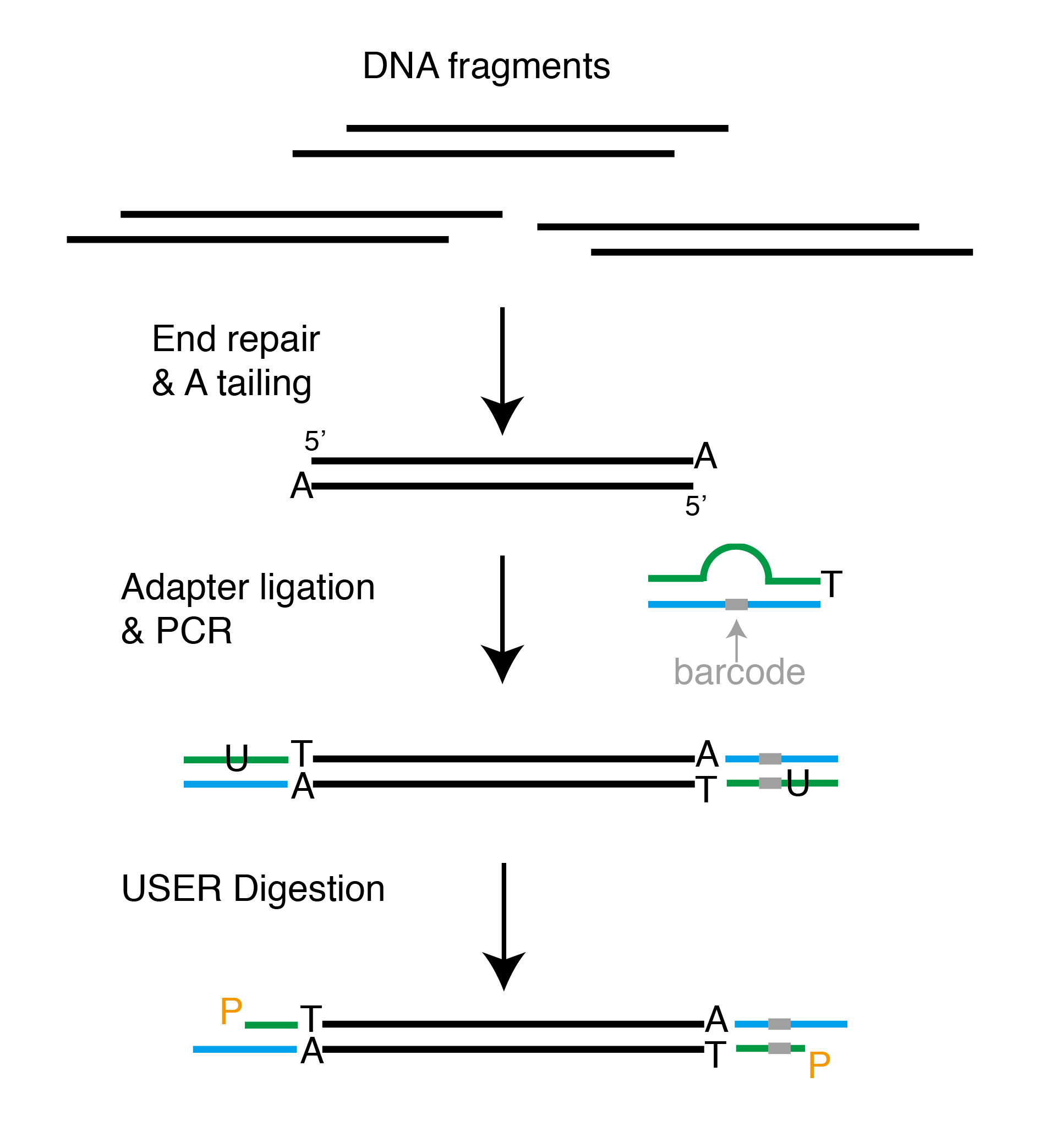

**Supplementary Figure S2. Scheme Presentation of Ligation with Bubble Adapters.** DNA fragments are end-repaired and A tailed prior to adapter ligation. Bubble adapter is indicated as a fraction of adapter sequence (green) is mismatching the other strand (blue), and is incorporating with U and T-tailed. The barcode sequence is shown by grey bar. After adapter ligation and USER digestion, the structure of the final product with adapter is shown in the bottom.

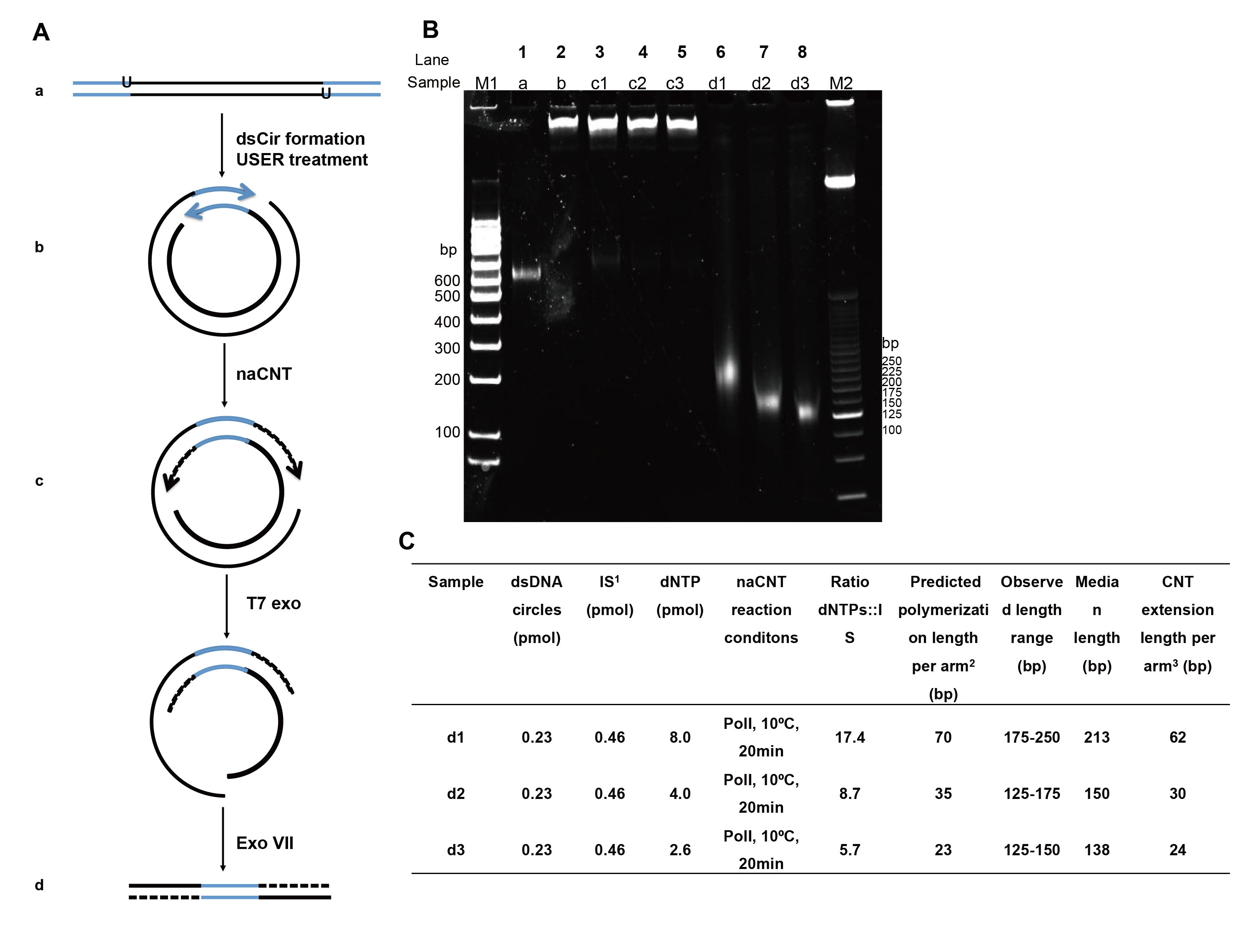

**Supplementary Figure S3. Evaluation of Controllable Nick Translation by Limiting Nucleotide Amount (naCNT).** (A) Schematic representation of library construction using naCNT. Genomic DNA fragments [indicated by black lines, sub-figure (a)] with directional adapters (blue lines) were treated with USER enzyme to create a 14-bp complementary overhang double-stranded DNA circles (dsCirs) were formed with two extendable gaps located 89-bp apart on opposite strands. After circularization and linear DNA digestion, naCNT was performed on the dsCirs [dashed black lines, figure (b)]. Then, DNA products were treated with T7 Exonuclease and Exonuclease VII (ExoVII) to remove nucleotides from the 5’-ends of dsDNA and degrade single-stranded DNA (c). The resulting products contain the adapter flanked by DNA templates created from the nick translation reaction (d). (B) Size analysis of products during different steps of the naCNT process using Pol I with 0.23 pmol of dsCirs and different amounts of dNTPs at 10°C for 20 min. Products of steps a-d [see figure (A)] were run on 6% polyacrylamide gels. Conditions 1 to 3 are depicted in the table in (C). Markers M1 and M2 represent ThermoFisher MassRuler Low Range DNA Ladder and 25 bp DNA Ladder, respectively. (C) Summary table of the detailed experimental conditions and calculations for each sample: ^1^The number of initiation sites (ISs) is two times the quantity of dsDNA circles used (pmol). ^2^The predicted polymerization length for each arm is equal to the ratio of dNTPs to gap sites multiplied by 4. ^3^The calculated polymerization length for each arm is equal to the median observed length minus the adapter length (90 bp) and all divided by 2 (for CNT and CPE region, respectively).

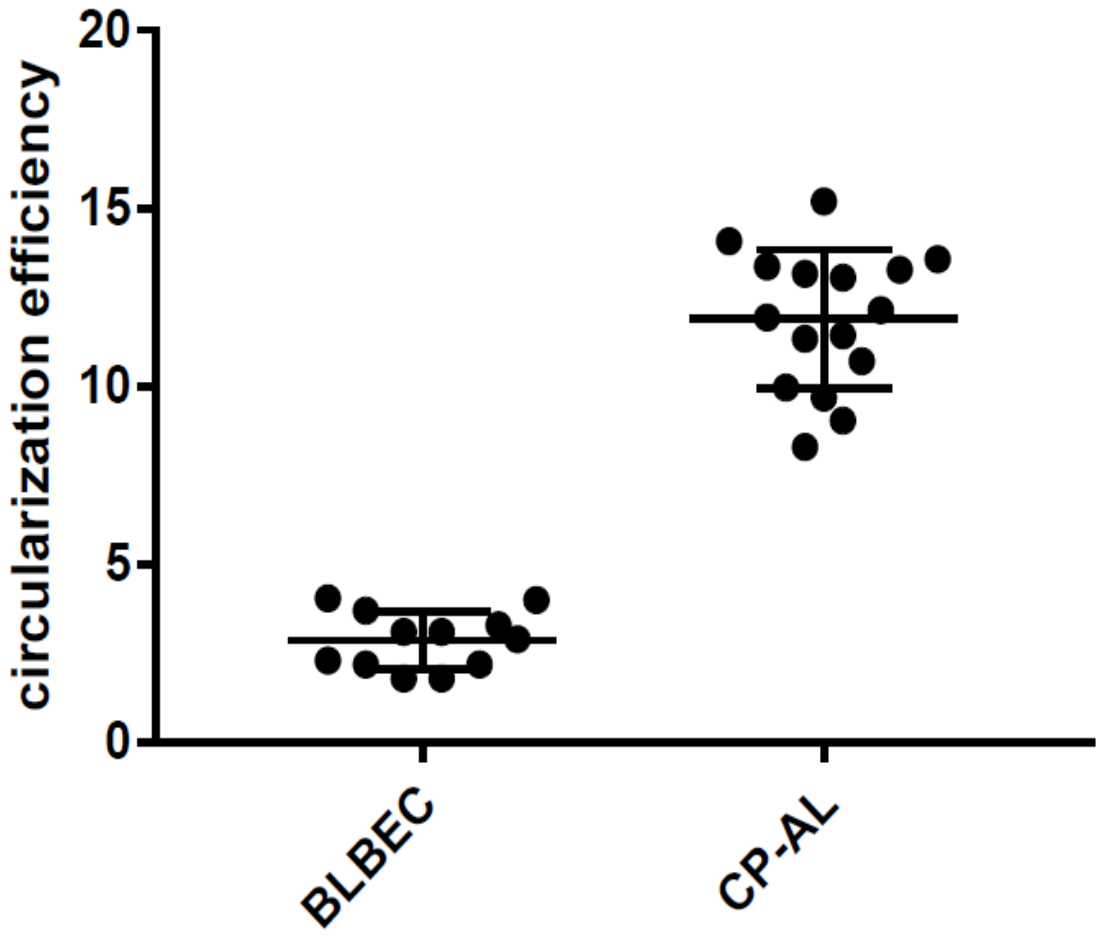

**Supplementary Figure S4. Evaluation of Circularization Efficiency between Two Methods.** Boxes show 16 replicates with our method (CP-AL), while circles indicate 12 samples with BLBEC (biotin-labeled blunt-end circularization) method (non-size-selected). Wilcoxon rank-sum test shows a significant difference between the two groups (P<0.0001), and by comparing the median value, CP-AL (12%) has over 4-fold change than BLBEC method (3%). Circularization efficiency was calculated as the ratio of the DNA amount remaining (after linear DNA digestion) to the original DNA input.

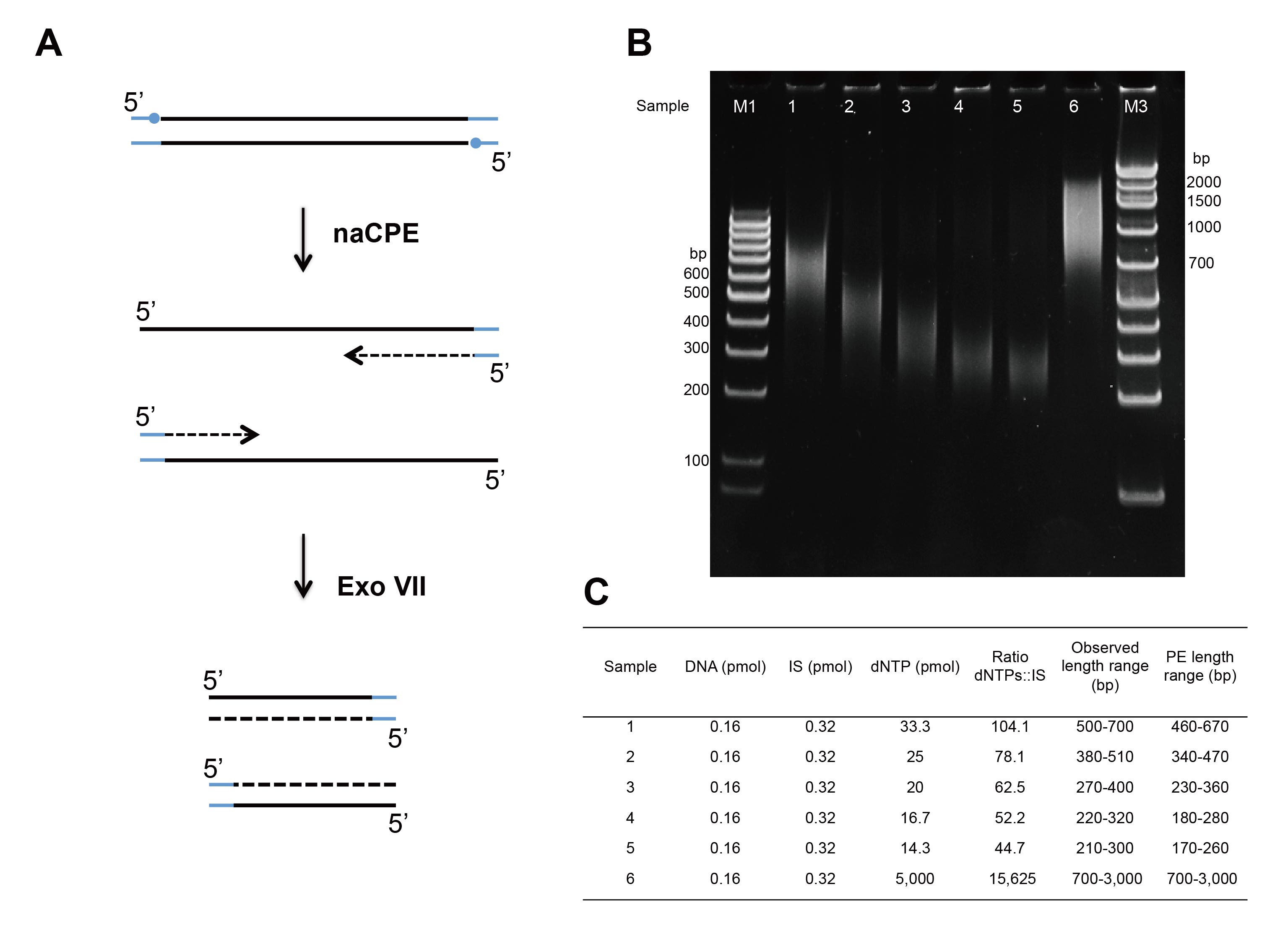

**Supplementary Figure S5. Evaluation of Controllable Primer Extension by Limiting Nucleotide Quantities (naCPE).** (A) Schematic representation of naCPE reaction. 3’-dideoxy-protected blunt-end adapters (indicated by blue lines with solid circles) were ligated to DNA fragments (black lines) ranging from 700 bp to 3 kb. The ligated products were then denatured, annealed with a CPE primer, and extended with Taq polymerase and titrated quantities of dNTPs. After the naCPE reactions, ExoVII treatment was used to degrade the 5’ single-stranded overhangs. The newly synthesized strand is depicted using dashed lines. (B) Size analysis of products after naCPE using Taq Polymerase, 0.16 pmol of dsDNA template, and different quantities of dNTPs (see C). Products were run on a 6% polyacrylamide gel. M1 = ThermoFisher MassRuler Low Range DNA Ladder; M3 = GeneRuler 1kb Plus DNA Ladder. (C) Table of observed size ranges based on dNTP quantity.

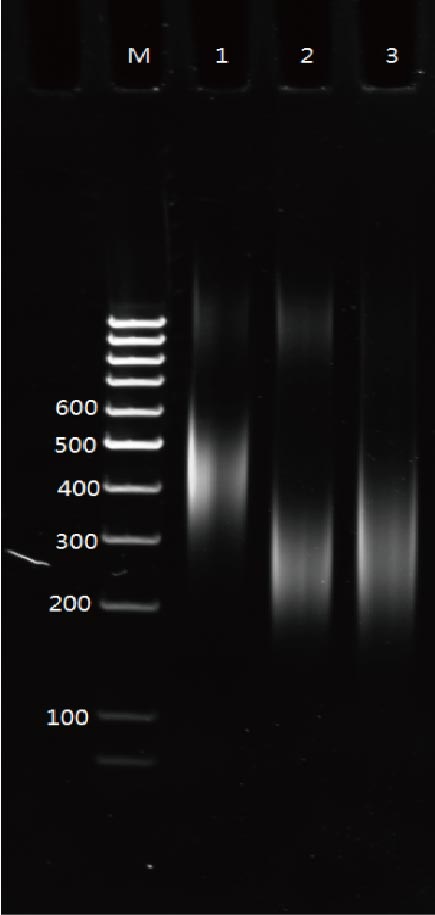

**Supplementary Figure S6. Gel analysis of the final products and pair-end regions.** Gel figure shows the size distribution of PCR products: Lane 1: full length of final products amplified by the primers annealing to Ad2_5’ and Ad2_3’ (referring to **Figure 1**); Lane 2: Paired-end region amplified by the primers annealing to Ad1 and Ad2_3’; Lane 3: Paired-end region amplified by primers annealing to Ad1 and Ad2_5’. Since the PCR products from lane 2 and 3 include approximately 50 bp of primer sequences, the minimal sizes of both pair-end regions are 150-bp, respectively.

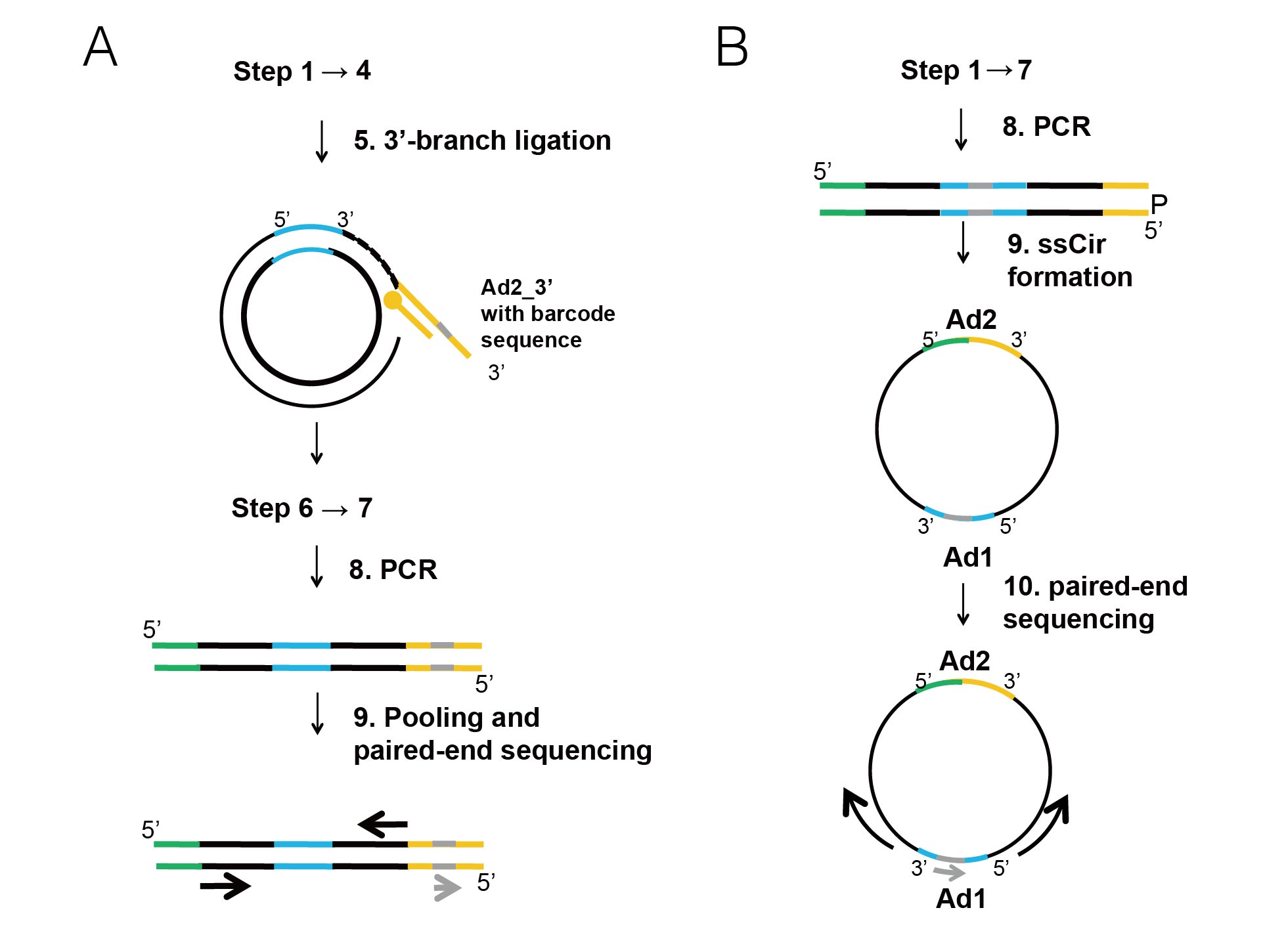

**Supplementary Figure S7. Paired-end indexed sequencing method for linear double strand DNA and single strand DNA.** (A) Scheme representation of linear double strand DNA index library construction. After 3’branch ligation, the sample barcode sequence (grey bar) is added through Ad2_3’ indexed adapter ligation prior to controllable primer extension. After primer extension reaction, individual libraries with different barcode sequences will be pooled with equimolar quantities as one library after Ad2 PCR and subjected for paired-end sequencing. DNA template, Ad1, Ad2_5’ and Ad2_3’ are indicated in black bar, blue, green and yellow, respectively while the sequencing orientations for the paired-end reads and the barcode sequences are indicated by two black arrows and a red arrow, respectively. (B) Scheme representation of single strand DNA index library construction. The barcode sequence is incorporating in Ad1 (grey bar), and after single strand circle formation, the sequencing orientations can be starting from bi-directions of Ad1 (indicated by black arrows) with barcode information sequencing (grey arrow).

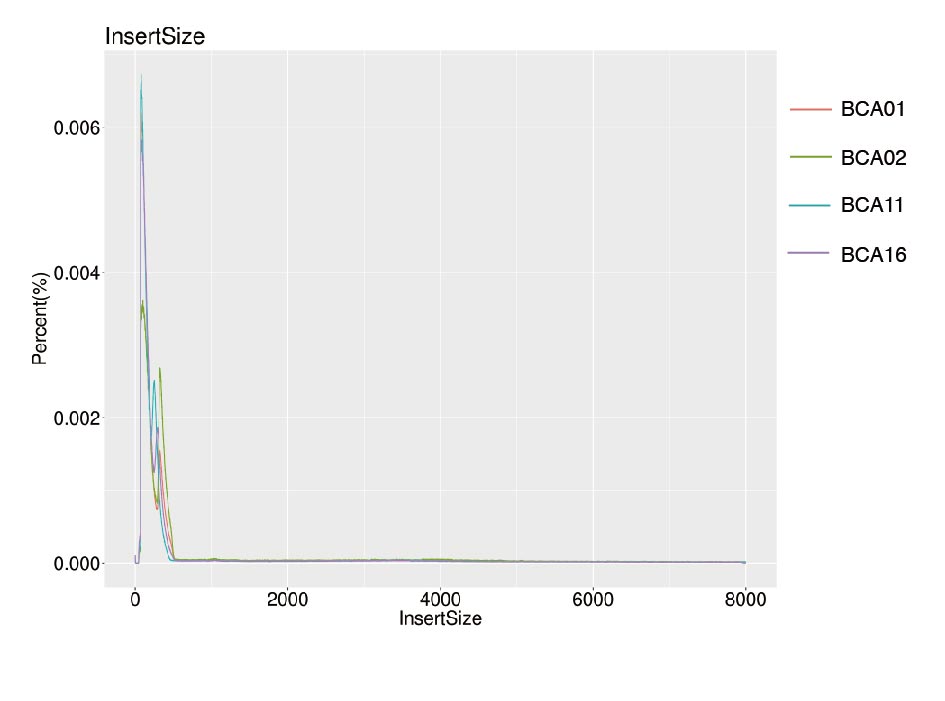

**Supplementary Figure S8. Distribution of “inward” read-pairs in WGS data with BLBEC library construction.** “Inward” read-pairs in four cases with BLBEC libraries are defined as read-pairs aligned to the genome with *cis*-orientation (+) in smaller coordinate while with trans-orientation (-) in larger coordinate. The percentage of “inward” read-pairs among the total read-pairs in these four samples are 27.5% while the ~91.6% of them are with an aligned distance <1-kb.

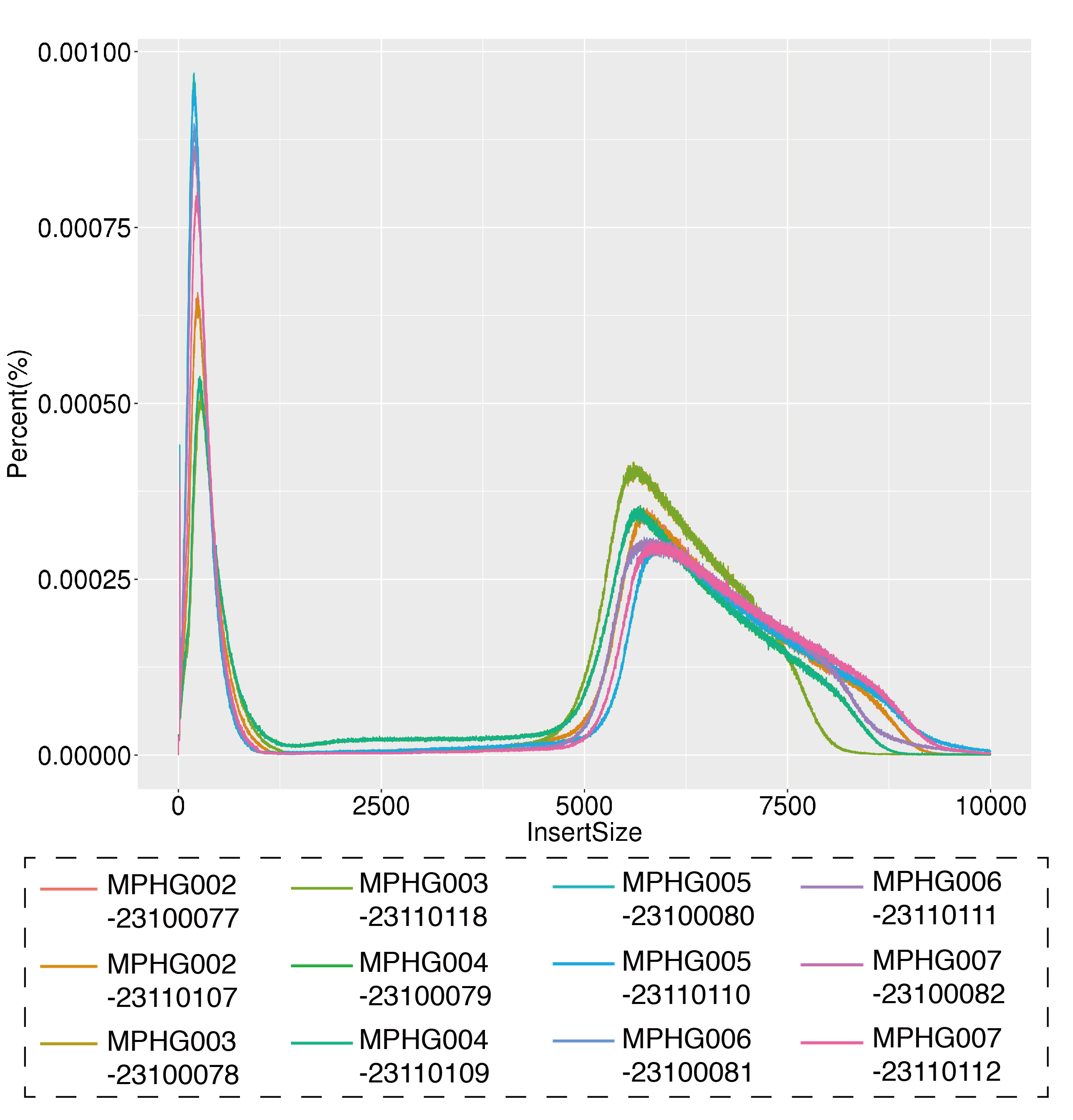

**Supplementary Figure S9. Evaluation of Insert-size Distribution in Data from Nextera.** The size distribution of 12 datasets of six samples with 6-kb insert size (gel-selected and read length trimmed as 100 bp) Nextera Mate-pair Library Construction Kit.

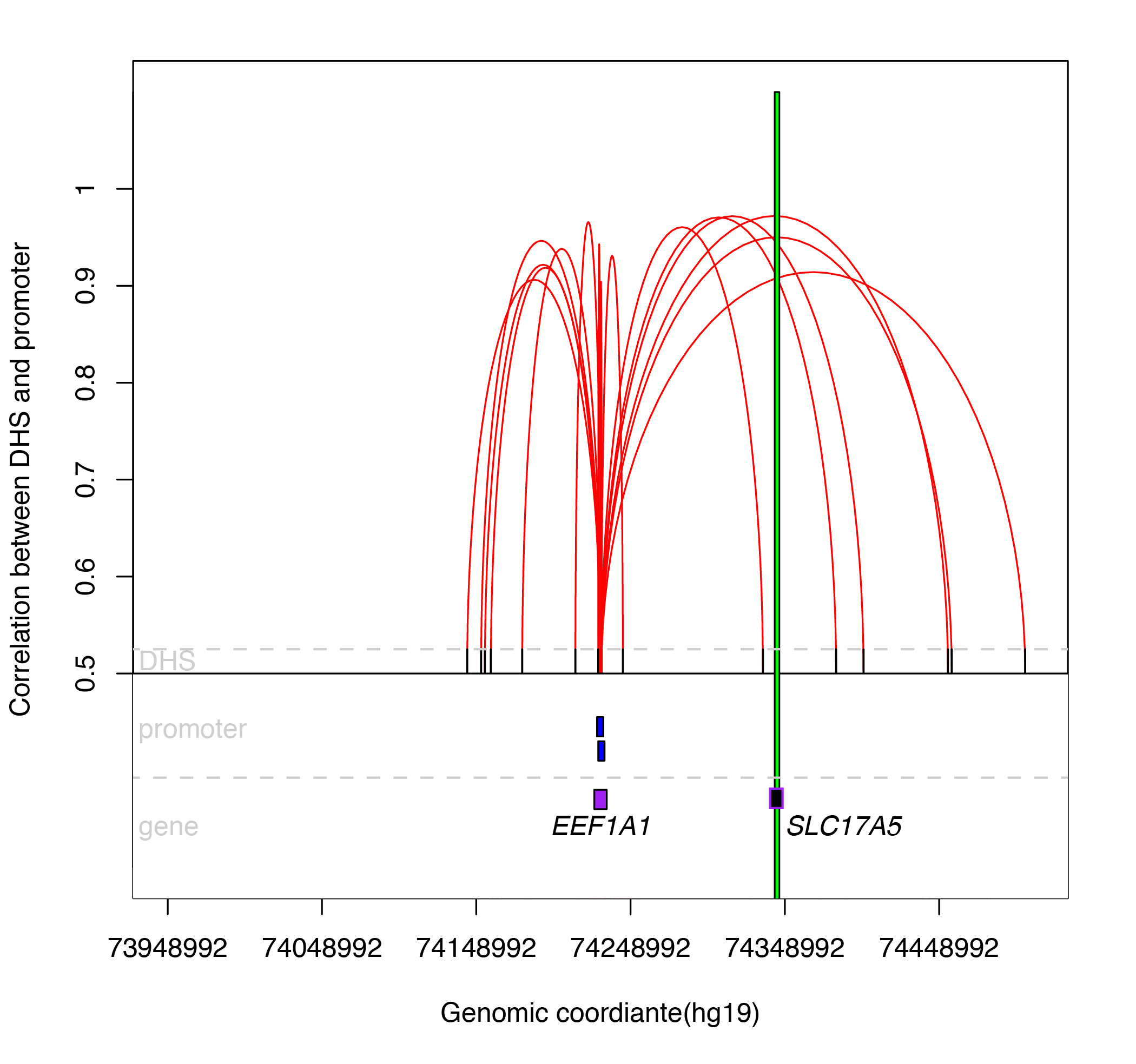

**Supplementary Figure S10. Disrupting Interactions between DNa I Hypersensitive Sites (DHSs) and Promoters of *EEF1A1* by the Breakpoint.** Figure indicates the diagram of the cross-cell-type correlation between distal DHSs and promoters of gene *EEF1A1* based on the reported map^4^. X axis represents the genomic coordinate of each element (such as gene and promoter), while Y axis shows the value of each correlation (r >0.9, reflected by a red line) between distal DHSs (indicated by black bar) and promoters (blue bar) of gene *EEF1A1* (purple box). This figure indicates there are 33.3% (5/15) cross-cell-type correlations between distal DHSs and promoters of gene *EEF1A1* are disrupted by the breakpoint (a green vertical line), which disrupted gene *SLC17A5* (blue box).

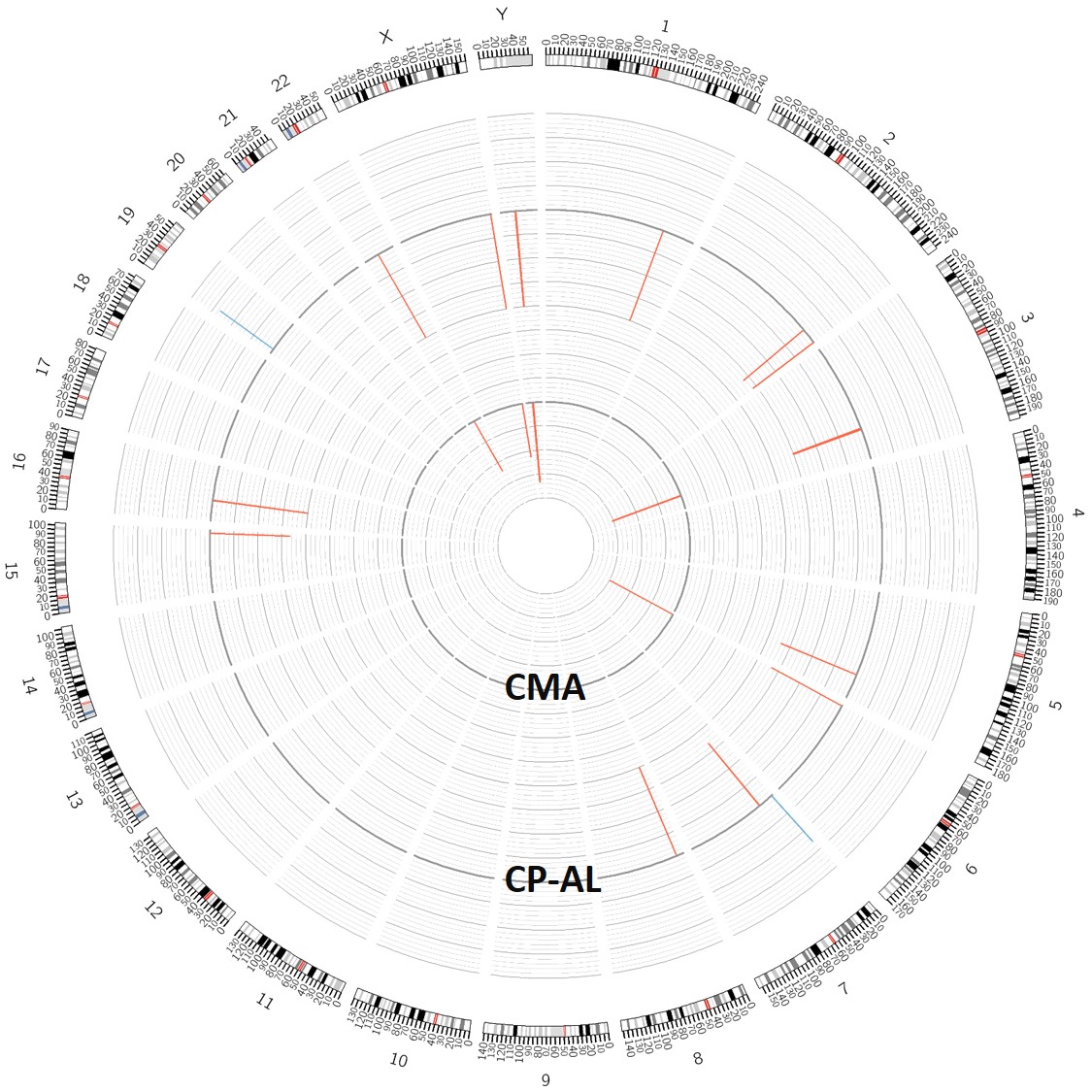

**Supplementary Figure S11. CNV detection between CP-AL and CMA.** The distribution of CNVs by CMA (inner circle) and CP-AL (outer circle) in the samples from six patients. Karyotypical structures and cytogenetic band colors are shown according to the University of California, Santa Cruz Genome Viewer Table Browser and chromosome color schemes (outmost circle). Rectangles in red and blue indicate consistent copy-number losses and gains, respectively, detected by both methods.

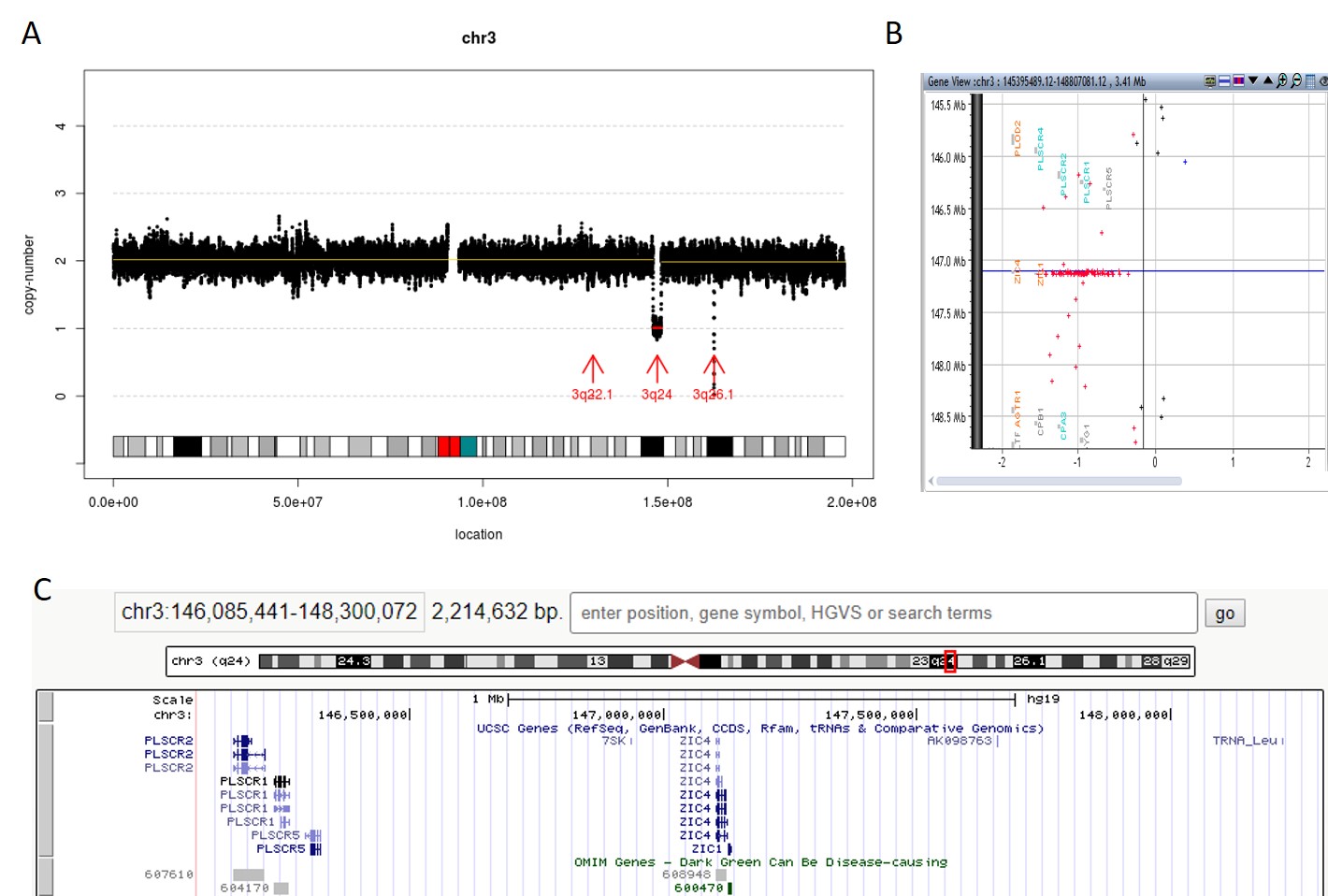

**Supplementary Figure S12. 3q24 deletion detected in Sample05**. (A) 2.2Mb deletion detected in 3q24 (indicated by a red horizontal line with a red arrow labelled as 3q24) by CP-AL. Each dot in black indicates an adjustable sliding window (50-kb with 5-kb increment). X axis shows the genomic location of each window, while Y axis indicates the copy-number. (B) Consistent finding of 3q24 deletion from CMA result. Each dot represents a probe in CMA platform. Dot in red or blue indicates a copy-number loss or a copy-number gain in case sample compared with control, respectively, while dot in black represents a copy-number neutral region. (C) The deleted region reported by CP-AL shown in UCSC genome browser with RefSeq genes and OMIM genes highlighted. *ZIC1* is the OMIM disease-causing gene within this region (highlighted in dark green).

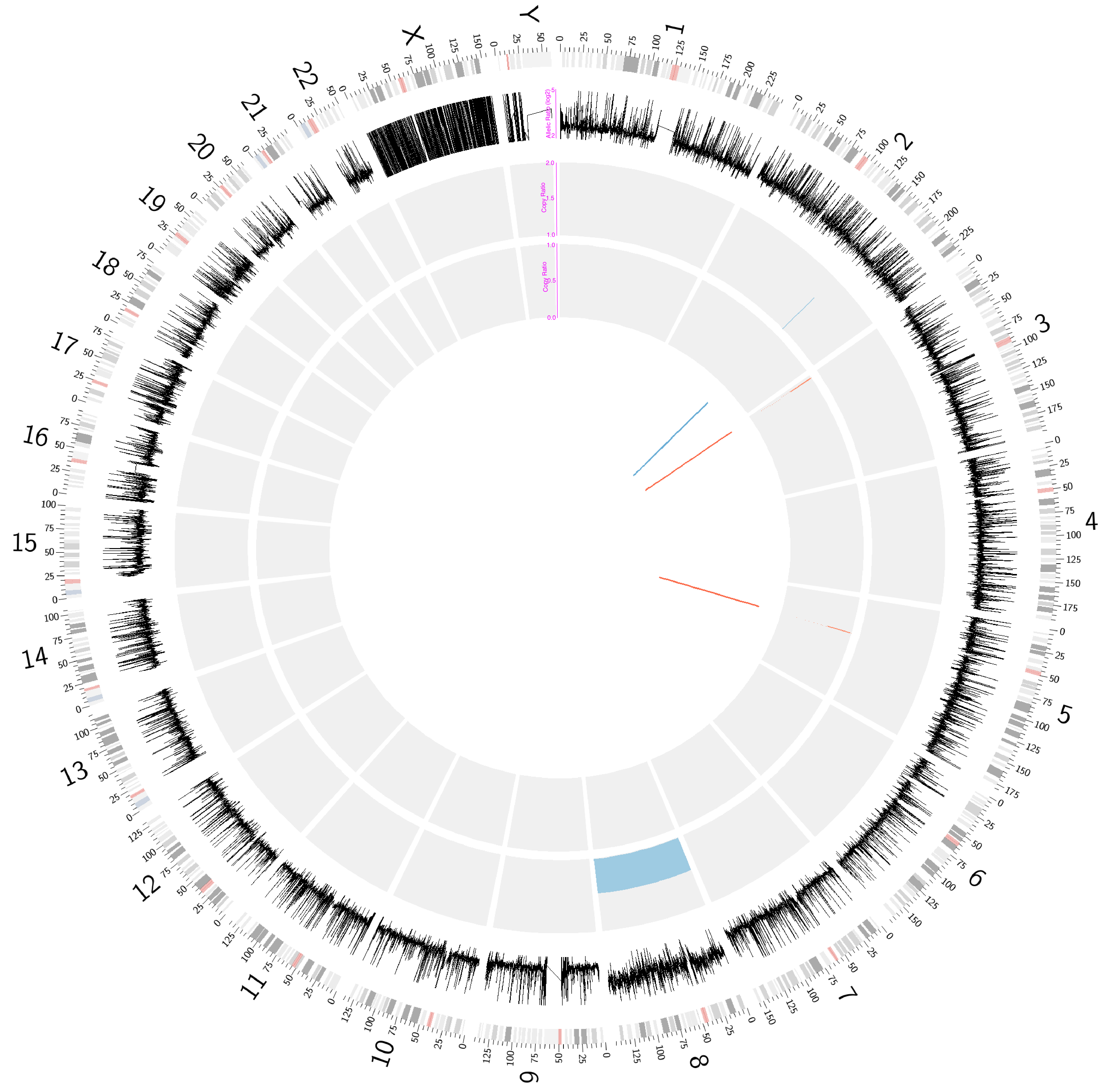

**Supplementary Figure S13. Whole-genome Analysis of Genomic Variants in Sample with Trisomy 8.** The distributions of allelic ratio (window-size: 100-kb in log2 scale; **Supplementary Methods**), copy-ratio and structural variants are shown from outer circle to inner accordingly. Allelic ratio of on average 4 is shown in chromosome 8 indicating there are two copies of one base type against only one copy of another base type. Karyotypical structures and cytogenetic band colors are shown according to the University of California, Santa Cruz Genome Viewer Table Browser and chromosome color schemes (outmost circle). Rectangles in red and blue indicate copy-number losses and gains, respectively, while lines in red and blue also indicate copy-number losses and grains.

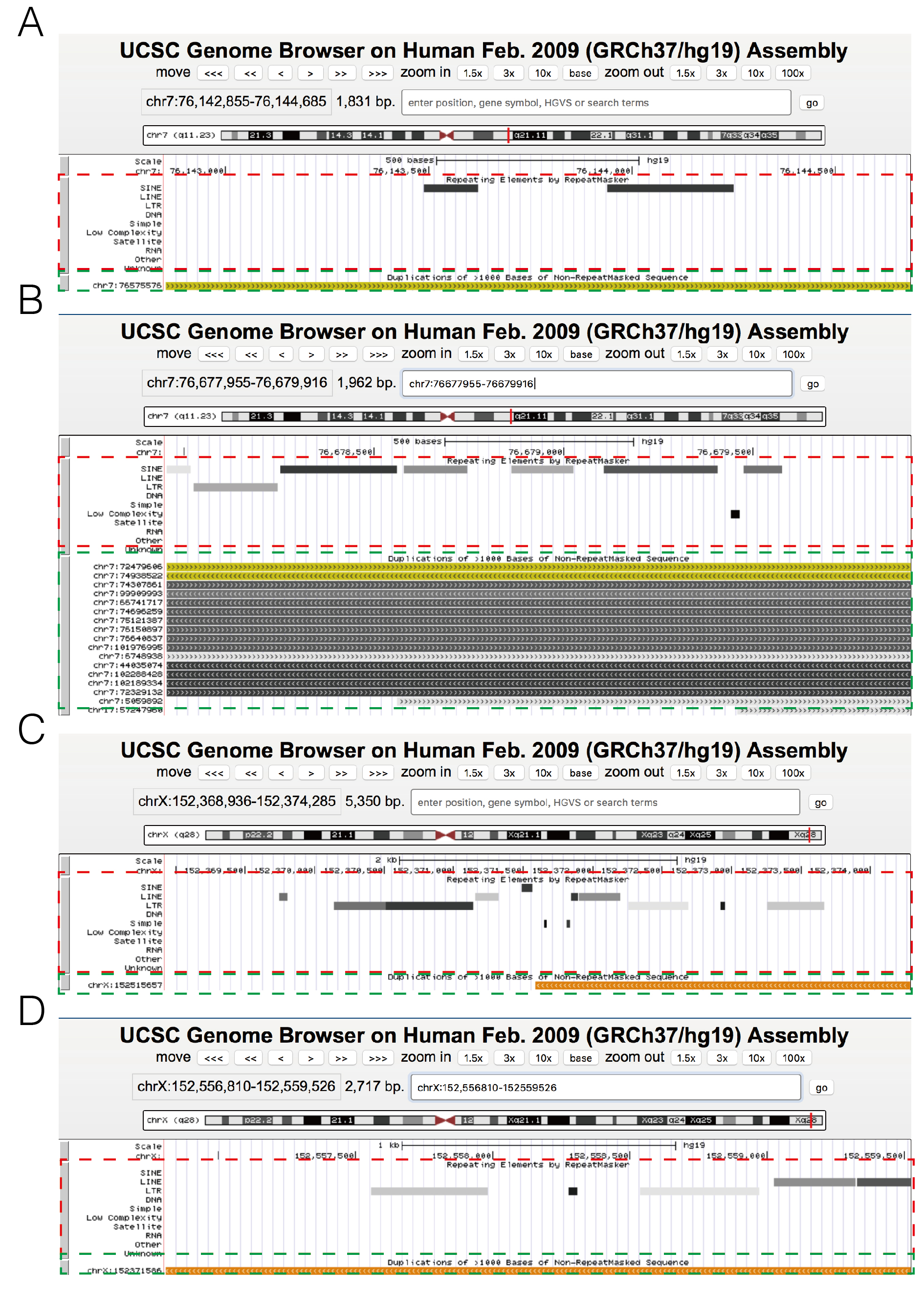

**Supplementary Figure S14. Distribution of Repetitive Elements in the Breakpoint Regions Covered by Read-Pairs of SV Inversions.** Figure (A) and (B) are the breakpoints regions indicated in the 533.2 kb cis-duplication detected in chromosome 6q14.1, while figure (C) and (D) are from the 187,9 kb inversion observed in chromosome Xq28. In each figure, repetitive elements were shown in a red dotted frame and segmental duplications were indicated in a green dotted frame. The figures were extracted from the information provided from the University of California, Santa Cruz Genome Browser (hg19).

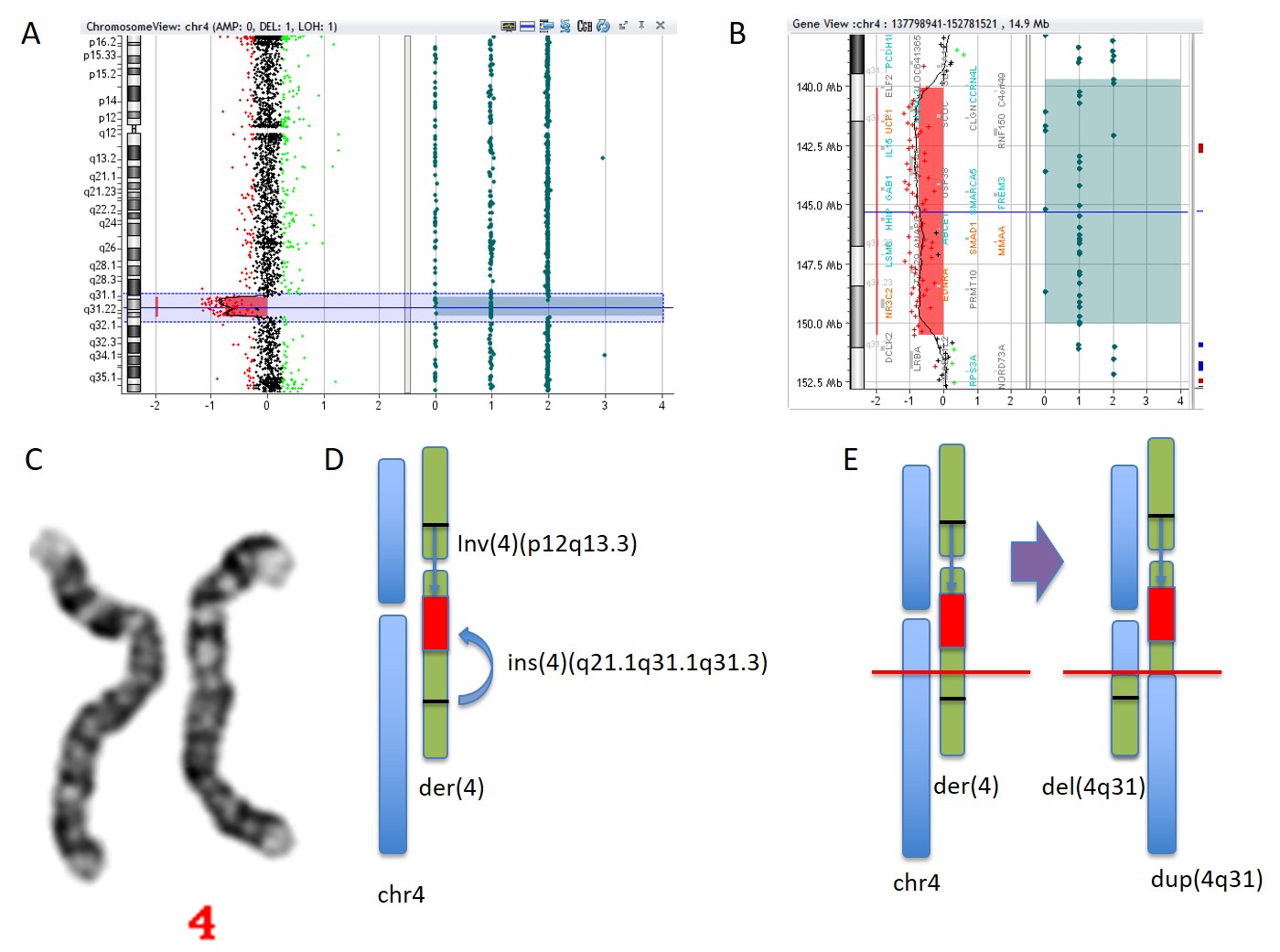

**Supplementary Figure S15.** Interstitial Deletion Detected Resulted from Paternal Intra-chromosomal Insertion. Figures (A) and (B) identified an interstitial deletion arr[hg19] 4q31.1q31.23(140046328_150534134)x1 approximately 10-Mb in size in the POC specimen. Each dot represents a probe in CMA platform. Dot in red or green indicates a copy-number loss or a copy-number gain in case sample compared with control, respectively, while dot in black represents a copy-number neutral region. (C) Karyotype in the paternal sample (Sample04) shows a derivative chromosome 4 with complex rearrangements. (D) The rearrangement events indicated by CP-AL. Blue bars show the normal chromosome 4, while green bars indicate the derivative chromosome 4. Red bar represents the deleted segment (4q31.1q31.23) and inserted to 4q21.1 (indicated by a blue curved arrow) involving an inversion inv(4)(p12q13.3). (E) Suspected mechanism of causing deletion in POC during crossing-over of the meiosis I during sperm genesis. The normal and derivative chromosomes 4 were suspected to be pairing and recombination occurred in a certain region (indicated by a red horizontal line) resulting in a chromosome 4 with 4q31.1q31.23 deleted and a chromosome 4 with 4q31.1q31.23 insertion and inversion involved.

**Supplementary Table S1. Data Demographic of Two Cases with Trisomy**

| Sample ID | Number of read-pairs (million) | Read-depth (fold) | Mapping rate (%) | Unique rate (%) | Mismatch rate (%) | Coverage (%)^1^ |
| --- | --- | --- | --- | --- | --- | --- |
| Trisomy 2 | 484 | 32.3 | 99.09 | 96.50 | 0.47 | 99.18 |
| Trisomy 8 | 544 | 36.3 | 98.69 | 96.68 | 0.44 | 99.41 |

^1^ Percentage of coverage was calculated as the number of bases covered with no less than N-fold dividing by the non-N region size of the human genome reference (hg19).

**Supplementary Table S2. Workflow and Estimated Turn-around-time for CP-AL**

| **Steps** | **Description** | **Estimated time** |
| --- | --- | --- |
| **1** | **DNA preparation and qualification** | Varied by the  laboratory  practice |
|  | Genomics DNA extraction |  |
|  | DNA quantitation and QC |  |
| **2** | **DNA fragmentation (HydroShear)** | ~1.5h |
|  | Shear the DNA |  |
|  | Fragment size QC (via agarose gel ,optional) |  |
|  | Purification |  |
|  | DNA quantitation |  |
| **3** | **End Repair** | ~1h |
|  | Purification |  |
| **4** | **A-tailing** | ~1h |
|  | Purification |  |
| **5** | **Adapter ligation (Ad1)** | ~1h |
|  | Purification |  |
|  | DNA quantitation |  |
| **6** | **PCR amplification (Ad1)** | ~5h  (Day1 stop point) |
|  | Purification |  |
|  | DNA quantitation |  |
| **7** | **dsDNA circularization** | ~6h |
|  | USER treatment |  |
|  | dsDNA circularization |  |
|  | Purification |  |
|  | Linear DNA digestion |  |
|  | Purification |  |
|  | DNA quantitation |  |
| **8** | **CNT & 3’-branch ligation** | ~2h  (Day2 stop point) |
|  | CNT |  |
|  | 3’-branch ligation |  |
|  | Purification |  |
| **9** | **CPE & 3’-branch ligation** | ~2.5h |
|  | CPE |  |
|  | Purification |  |
|  | 3’-branch ligation |  |
|  | Purification |  |
| **10** | **PCR amplification (Ad2)** | ~1.5h  (Day3 stop point) |
|  | Purification |  |
|  | DNA quantitation |  |
| **11** | **Preparation for sequencing** | Depends on platform |

**Supplementary Table 3. Sensitivity and specificity of SNV and InDel detection**

| Library | Nick translation | Primer extension^1^ | SNV detection (%)^2^ | InDel detection (%)^2^ |
| --- | --- | --- | --- | --- |
| #1 | PolI | ttCPE by PfuCx | 95.2\|98.3 | 80.7\|93.6 |
| #2 | PolI+2xAT^3^ | ttCPE by PfuCx | 96.1\|99.3 | 82.4\|94.5 |
| #3 | PolI+2xAT^3^ | naCPE by Taq | 97.3\|99.7 | 84.6\|90.6 |
| #4 | PolI/Taq+2xAT^3^ | naCPE by Taq | 96.0\|99.7 | 81.4\|92.8 |

^1^ ttCPE and naCPE refer to controlled primer extension by adjusting of reaction temperature and duration time or limiting the nucleotide input only

^2^ Sensitivity and specificity of SNV and InDel detection were calculated by comparing the detection results from CP-AL with those reported by using small-insert libraries sequenced in the same platform .

^3^ Additional two-fold of dATP and dTTP relative to dGTP and dCTP
